# Iron retention coupled with trade-offs in localized symbiotic effects confers tolerance to combined iron deficiency and drought in soybean

**DOI:** 10.1101/2025.01.02.631154

**Authors:** Md Rokibul Hasan, Asha Thapa, Ahmad H. Kabir

## Abstract

Iron (Fe) and water availability are closely interlinked, with deficiencies in both adversely affecting soybean growth. However, the strategies employed by soybean to tolerate such conditions remain poorly understood. This study elucidates the interactions of host factors, and microbial associations using multi-omics approaches in Clark (tolerant) and Arisoy (sensitive) genotypes exposed to Fe deficiency and drought. Clark exhibited resilience to stress through sustained osmotic regulation, nutrient uptake, and photosynthetic activity, in contrast to Arisoy. Particularly, Fe retention in Clark, accompanied by the upregulation of ferritin-like proteins, may mitigate oxidative stress by reducing Fenton reactions. Furthermore, higher jasmonic and salicylic acid levels in Clark may contribute to its enhanced stress adaptation compared to Arisoy. RNA-seq analysis revealed 818 and 500 upregulated, along with 931 and 361 downregulated genes, in the roots of Clark and Arisoy, respectively, under stress. We observed the upregulation of symbiotic genes, such as *Chalcone-flavonone isomerase* 1 and *SWEET10*, accompanied by increased rhizosphere siderophore and root flavonoid in Clark. This indicates a significant role of microbes in mediating differential stress tolerance in soybean. Particularly, the combined stress led to distinct root and nodule microbiome dynamics, with Clark recruiting beneficial microbes such as *Variovorax* and *Paecilomyces*, whereas Arisoy exhibited the opposite pattern. In addition, Clark maintained nodule *Bradyrhizobium* and tissue nitrogen status, supported by ammonium retention and induction of *Ammonium transporter* 1 in the roots. Furthermore, *in vitro* compatibility between *V. paradoxus* and *P. lilacinus* suggests a synergistic interaction, with their localized signals benefiting Clark. Remarkably, enriched microbiomes significantly improved growth parameters, accompanied by elevated rhizosphere siderophore in sensitive genotypes under stress. This study is the first to uncover mechanisms of dual stress tolerance in soybean that may offer promising targets for breeding programs and microbiome-based biofertilizer strategies to improve combined stress tolerance in soybean and other legumes.

**Highlight:** Iron retention coupled with symbiotic associations driven by the enrichment of *Variovorax* and *Paecilomyces* in the roots confers tolerance to combined iron deficiency and drought in soybean.

## 1. Introduction

Climate change poses an increasing threat to global agricultural sustainability by intensifying abiotic stresses, including drought and nutrient deficiencies. Drought significantly reduces crop growth and productivity by impairing critical physiological processes (Qiao *et al.*, 2024). This stress arises from insufficient soil moisture, which not only restricts water availability but also limits the uptake of essential minerals by plant roots (Li *et al.*, 2023). Iron (Fe) deficiency, a prevalent nutritional disorder in plants, results from low bioavailability in alkaline or calcareous soils, where high pH restricts Fe solubility and uptake (Li and Lan, 2017; Kabir *et al.*, 2012). Fe deficiency not only affects plant health but also disrupts photosynthetic efficiency, gene expression, and metabolic pathways (Liang, 2022), leading to symptoms like interveinal chlorosis in young leaves (Li *et al.*, 2021). Drought further exacerbates this condition by lowering soil moisture and Fe solubility, which is particularly critical in legumes, where Fe plays a key role in nitrogen fixation (Lindström and Mousavi, 2020; Briat *et al.*, 2015). Drought disrupts nutrient uptake and increases reactive oxygen species (ROS), thus hindering plant growth and survival (Hussain *et al.*, 2018). Therefore, understanding the interplay between Fe deficiency and drought responses in legume crops is crucial for devising strategies to improve resilience under complex stress conditions.

Plants utilize two main strategies to absorb Fe from the soil: Strategy I (non-graminaceous species) and Strategy II (graminaceous species) (Liang, 2022; Gill and Tuteja, 2010). Plants using Strategy I possess several mechanisms to enhance Fe uptake under Fe deficiency. One such adaptation involves increasing proton extrusion through *ATPase 1*, which acidifies the rhizosphere and improves Fe solubility (Kabir *et al.*, 2012). Additionally, ferric-chelate reductase (FCR) enzymes convert Fe(III) into the more bioavailable Fe(II) form (Kobayashi and Nishizawa, 2012; Morrissey and Guerinot, 2009). Plants further upregulate Fe transporter proteins to facilitate the uptake of bioavailable Fe into root cells (Kroh and Pilon, 2019). Plants adapt to drought by maintaining cellular hydration and minimizing damage through strategies such as modifying root architecture to enhance water uptake and closing stomata to reduce transpiration (Kumar *et al.*, 2021; D’Oria *et al.*, 2022). Furthermore, the ammonium transporter located on the plasma membrane actively transports ammonium ions from nitrogen-fixing bacteria in nodules of legume crops into the plant cell (Hao *et al.*, 2020). At the molecular level, plants undergo dynamic changes involving the activation of stress-responsive genes and signaling pathways. Several genes play a direct role in stress mitigation, including those encoding dehydrins (Hassan *et al.*, 2015) and enzymes involved in osmolyte biosynthesis (Ghosh *et al.*, 2021), which contribute to water retention. Furthermore, abscisic acid (ABA) signaling plays a central role by regulating stomatal closure, conserving water, and activating a cascade of stress-responsive genes under drought conditions (Chaves *et al.*, 2003; Ali *et al.*, 2020). Jasmonic acid (JA) protects plants by regulating stomatal closure by boosting antioxidant defenses and promoting osmotic adjustment to reduce cellular damage (Wasternack and Song, 2017; Dhakarey *et al.*, 2016). In drought-stressed plants, JA coordinates with other hormones like ABA and salicylic acid (SA) to regulate water loss and activate stress-responsive genes (Rai *et al.*, 2024; Yang *et al.*, 2019). However, the physiological strategies and gene activation mechanisms triggered in plants exposed to combined Fe deficiency and drought are not yet well understood.

While plants possess intrinsic mechanisms to mitigate abiotic stress, these are often insufficient without the support of beneficial microbes. Sugars and secondary metabolites such as flavonoids play crucial roles as host determinants influencing microbial recruitment, particularly under stress conditions (Loo *et al.*, 2024; Lidoy *et al.*, 2023). These compounds act as chemical signals in the rhizosphere, shaping microbial communities and promoting beneficial interactions that enhance plant resilience. The shifts in the belowground microbiome underlying abiotic stresses in plants involve changes in both the relative abundance and composition of microbial groups (Trivedi *et al.*, 2022). Under Fe-limiting conditions, rhizosphere microbes, including bacteria (*Pseudomonas, Variovorax*) and fungi (*Trichoderma*), secrete siderophores that solubilize Fe(III) for plant uptake (Kabir and Bennetzen, 2024; Wang *et al.*, 2024; Crowley *et al.*, 2006). A recent study revealed the enrichment of *Variovorax*, Shinella, and *Chaetomium* in the roots of pea plants under alkaline stress (Thapa *et al.*, 2025a). Drought significantly influences not only microbial diversity but also the composition of microbial communities in the rhizosphere and plant parts (Salamon *et al.*, 2020; Naylor and Coleman-Derr 2018). However, the dynamics of microbial communities and their synergistic relationships in response to combined stresses in legumes require further research.

Soybean (*Glycine max*), one of the most important legume crops globally, plays a vital role in food, feed, and biofuel production (Longley *et al.*, 2020). Its productivity is highly susceptible to Fe deficiency (Zocchi *et al.*, 2017) and drought (Du *et al.*, 2024), with their interactive effects more severely impairing key physiological processes and further reducing yield. Despite advances in understanding individual stress responses, the interplay of molecular pathways and microbial dynamics under combined Fe deficiency and drought remains unexplored in soybean. This study addresses this gap by integrating transcriptomic, physiological, and microbiome analyses to uncover synergistic mechanisms underlying combined stress resilience in soybean. Furthermore, we investigated the interactions and persistence of enriched microbial consortia and their synergistic interactions that induce resilience in sensitive genotypes of soybeans to Fe deficiency and drought. This novel study not only sheds light on the candidate genes and dynamics of soybean-microbe interactions under dual stresses but also lays the foundation for innovative strategies to enhance soybean resilience in the face of combined Fe deficiency and drought stress in the soil.

## 2. Materials and methods

### Plant cultivation technique

The seeds of contrasting genotypes of soybean (tolerant: Clark, sensitive: Arisoy, AG58XF3) to the combined Fe deficiency and drought were collected from USDA-GRIN and LSU AgCenter. Seeds were surface sterilized by immersing them in a 4% sodium hypochlorite solution for 5 min, followed by three rinses with sterile water, and then germinated in a tray. After 3 d, uniform, and healthy seedlings were transplanted into 500 g soil pots. The pots contained a 1:2 mixture of natural soil (soybean field) and commercial potting mix [peat moss, bark, and perlite (2:2:1)] without additional fertilizers and were used for two treatments: control (soil without lime, pH ∼6.5) and Fe-drought+ (soil amended with 4.5g NaHCO_3_ and 3.0g CaCO_3_, pH ∼7.8). The physicochemical characteristics of the soil are detailed in Supplementary Table S1. During the cultivation period, NaHCO_3_ was added weekly to maintain the soil pH at 7.8 making Fe unavailable to plants (Kabir *et al.*, 2012). In addition, drought conditions were induced two weeks after transplanting. Soil moisture content was monitored daily to maintain water levels at 75% for control plants and 25% for drought-stressed plants. To study the impact of microbial absence on plant stress responses, autoclaved vermiculite was used in pots for cultivation in a separate experiment. The induction of Fe deficiency and drought stress was performed as described previously; however, the amounts of NaHCO_3_ and CaCO_3_ were reduced to 3.0g and 2.0g, respectively, to maintain the pH at ∼7.8. For the microbial consortia study, *Variovorax paradoxus* and *Paecilomyces lilacinus* were inoculated into the soil at a concentration of 1×10^9^ CFU per gram. The plants were grown in a randomized complete block design with 9 replications (3 plants per pot) per treatment in the greenhouse at 25°C, with a photoperiod of 10 h light/14 h darkness and a light intensity of 250 μmol m^-2^ s^-1^. The plants were cultivated for 4 weeks before data collection.

### Determination of morphological and physiological parameters

A digital caliper was used to measure the height and length of the shoot and root, respectively. The dry weights of the roots and shoots were then measured after they had been dried for 3d at 75°C in an electric oven. Also, nodules were carefully collected from soybean roots by gently washing the roots to remove soil, followed by manual detachment of the nodules using sterilized forceps. A handheld chlorophyll meter (AMTAST, United States) and a FluorPen FP 110 (Photon Systems Instruments, Czech Republic) were used to measure the chlorophyll score and Fv/Fm (maximal photochemical efficiency of PSII) on the uppermost fully developed leaves at three distinct sites. The leaves were exposed to darkness for one hour before data collection to prepare them for OJIP analysis. Furthermore, the relative water content (RWC) of fully expanded trifoliate leaves was measured from the tip of the main stem (Schonfeld *et al.*, 1988). Briefly, the fresh weight (FW) of the leaves was recorded immediately after excision. After soaking the leaves in distilled water for a full day at room temperature, they were thoroughly blotted dry using tissue paper, and the turgid weight (TW) was determined. The leaf samples were oven-dried for 72 h at 75°C to estimate their dry weight (DW). RWC was calculated using the formula: RWC (%) = (FW – DW) / (TW – DW) x 100. Furthermore, young leaves and roots taken from plants were subjected to nutrient analysis. After being separated, the roots were rinsed with deionized water, immersed in a 0.1 mM CaSO_4_ solution for ten min, and then cleaned under running tap water. In addition, deionized water was used to wash the young leaves individually. After being wrapped in envelopes, each sample was dried for 3d at 75°C in an electric oven. Elemental concentrations were determined using inductively coupled plasma mass spectrometry (ICP-MS) at the Agricultural and Environmental Services Laboratories, University of Georgia, following standard protocols.

### Analysis of ROS and Fenton reaction rate

For H_2_O_2_ determination, root samples were processed as previously described (Alexieva *et al.*, (2001). Briefly, the root samples were mixed with a 0.1% trichloroacetic acid solution and finely ground into a powder using a mortar and pestle. The homogenized extract was centrifuged at 10,000 rpm for 15 min to remove cellular residues. The resulting supernatant was supplemented with 1 M potassium iodide and 10 mM phosphate buffer (pH 7.0), and the mixture was incubated in darkness for 60 min. The optical density of the solution was subsequently measured at 390 nm using a spectrophotometer (ThermoFisher, Waltham, USA). The concentration of H_2_O_2_ was quantified using a standard curve. Furthermore, hydroxyl radicals (•OH) in the roots were determined as previously described (Coudray and Favier 2000). Briefly, fresh root tissues were harvested, thoroughly washed with deionized water, and homogenized in ice-cold sodium phosphate buffer (50 mM, pH 7.4) containing 1 mM EDTA. The homogenate was centrifuged at 12,000 × g for 10 min at 4°C, and the supernatant was collected. The reaction was initiated by incubating the supernatant with 5 mM salicylic acid in sodium phosphate buffer at 37°C for 30 min in the dark, allowing for the formation of 2,3-DHBA (2,3-dihydroxybenzoic acid). The reaction was stopped by adding cold ethanol (95% v/v), followed by centrifugation to remove debris. The concentration of 2,3-DHBA in the supernatant was quantified spectrophotometrically by measuring absorbance at 510 nm. A standard curve prepared with known concentrations of 2,3-DHBA was used for quantification. Furthermore, the Fenton reaction rate for root samples was determined by the concentration of Fe and as H_2_O_2_ using the following formula: Fenton rate (µmol/s) = *k* × {Fe (µM)} × {H_2_O_2_ (µM), where *k* = 0.01 µM^-1^S^-1^ at pH 7 and 25°C.

### Ferric chelate reductase activity in roots

Ferric chelate reductase activity in roots was determined using sodium bathophenanthroline disulfonic acid (Na -BPDS) as the Fe^2+^-specific chelator (Waters *et al.*, 2002). Root tissues from the two genotypes were first rinsed thoroughly with deionized water, and root tips were cut and kept on ice. Approximately 100 mg of root tissue was washed in 0.2 mM CaSO_4_ solution for 5 min and subsequently transferred into 2 mL microcentrifuge tubes containing an assay solution consisting of 10 mM CaSO_4_, 5 mM MES buffer (pH 5.5), 0.1 mM Fe-EDTA, and 0.3 mM Na - BPDS. A separate tube containing the assay solution without root tissue served as the control. Both the sample and control tubes were incubated in a shaking water bath at 23°C for 1 hour at 3000 × g under dark conditions. Afterward, 1 mL of the assay solution was transferred to a cuvette, and the absorbance was recorded at 535 nm using a spectrophotometer. The ferric reductase activity was calculated based on the molar extinction coefficient of Fe(II)-BPDS (22.14 mM^-1^ cm^-1^).

### Phytohormone analysis in roots

The amounts of phytohormones in the roots were measured with Liquid Chromatography-Mass Spectrometry (LC-MS) as previously described (Zeng *et al.*, 2013). To put it briefly, 200 mg of fresh root tissue was crushed in liquid nitrogen and then extracted using an 80:20 (v/v) methanol: water solution that contained 0.1 g/L butylated hydroxytoluene and 0.1% formic acid. The samples were processed in a TissueLyser II (Qiagen, USA) at 30 Hz for 30 sec while being held in pre-frozen bead beater adapters. The samples were then incubated on a shaker at 5,000 rpm and 4°C for 16 h following a centrifugation for 20 sec at 4°C. The supernatant was then moved to autosampler vials for LC-MS analysis after they had been vortexed and centrifuged for 10 min at 4°C at 12,000 rpm. Each phytohormone standard (Sigma-Aldrich, USA) was made in a solution containing 0.1% formic acid and 80% methanol.

### RNA-seq analysis in roots

RNA-seq analysis was performed on root samples to investigate gene modulation under combined Fe deficiency and drought. The roots were carefully cleaned by washing twice with sterile ice-cold phosphate-buffered saline, including 10 sec of vortexing, followed by two rinses with ice-cold sterile water. The cleaned root tissues were ground into a fine powder using a pre-chilled mortar and pestle with liquid nitrogen. Total RNA was extracted from the powdered roots using the SV Total RNA Isolation System (Promega, USA). For RNA-seq library preparation, 1 μg of RNA with an RNA Integrity Number (RIN) above 8 was used. The libraries were prepared using the KAPA HiFi HotStart Library Amplification Kit (Kapa Biosystems, USA) and sequenced on an Illumina NovaSeq 6000 platform (PE 150) at the RTSF Genomics Core Facility, Michigan State University. Of the total reads generated, 92.5% passed quality control filters, with a quality score of ≥ Q30 (Supplementary Fig. S1).

Bioinformatics analysis of raw FastQ files was conducted using Partek Flow genomic analysis software (Partek, St. Louis, MO, USA). Adaptor sequences and low-quality reads were removed using Trimmomatic (Bolger *et al.*, 2014) to obtain high-quality clean reads. The clean reads were aligned to the soybean reference genome using HISAT2 (v2.0.5) (Kim *et al.*, 2015). Raw read counts were generated using HTSeq (Anders *et al.*, 2015) and imported into the iDEP 2.01 platform (Ge *et al.*, 2018) for differential expression analysis of genes (DEGs). The counts were normalized and log2-transformed using the voom method (Ritchie *et al.*, 2015) to account for library size and model mean–variance relationships prior to linear modeling in the limma package (v3.19). DEGs were defined as those with an adjusted *P*-value < 0.05 and a fold change ≥ 2, determined using the false discovery rate (FDR) method. MA plots for DEGs were created using ggplot2, and heatmaps were generated using pheatmap in the R platform. Gene enrichment analysis was performed using the ShinyGO 0.80, which provides functional annotation and pathway analysis specific to soybean genomic datasets (Ge *et al.*, 2020).

### Real-time qPCR validation of candidate genes

We performed real-time qPCR analysis to validate the expression of a few candidate genes, such as *GLYMA.20G241500* (*Chalcone-flavone isomerase 1*), *GLYMA.04G198400* (*Bidirectional sugar transporter SWEET10*), *GLYMA.19G091400* (Ferritin-like family protein), *GLYMA.10G146600* (Iron-regulated protein 3) and *GLYMA.17G027400* (Drought-induced-19 protein) in the roots of soybean on the CFX96 Real-time System (Bio-Rad, USA) with gene-specific primers (Supplementary Table S5). Briefly, the first-strand cDNA was synthesized from total RNA using the GoScript Reverse Transcription System (Promega, USA). The qPCR cycling conditions were as follows: initial denaturation at 95°C for 2 min, followed by 42 cycles of 95°C for 5 sec, annealing at 55°C for 30 sec, and a final extension at 60°C for 5 min. Gene expression was normalized using *GAPDH* and *Tubulin* as reference genes, whose Cq values showed minimal variability (1–1.5 cycles) across all samples, confirming their invariant expression under the experimental conditions. Relative expression levels were calculated using the 2^−ΔΔCt method (Livak and Schmittgen, 2001), with reference genes (*GAPDH* and *Tubulin*) serving as internal controls. The real-time PCR experiment included three independent biological replicates, with two technical replicates performed for each. The mean values of the biological replicates were used for statistical analysis.

### Determination of siderophore in the rhizosphere

Siderophore levels in rhizosphere soil were measured using the chrome azurol S (CAS) assay (Himpsl and Mobley 2019). Briefly, rhizosphere soil was collected by removing tightly adhering soil from the root surface. The soil was homogenized in 80% methanol and centrifuged at 10,000 rpm for 15 min. A 500 μl aliquot of the supernatant was mixed with an equal volume (500 μl) of CAS solution and incubated at room temperature for 5 min. Absorbance was measured at 630 nm, with a blank containing 1 ml of CAS reagent serving as the reference. The siderophore content per gram of soil was determined using the formula: % siderophore unit = [(A_ref - A_sample) / A_ref] × 100, where A_ref is the absorbance of the CAS reagent alone, and A_sample is the absorbance of the CAS reagent mixed with the soil sample.

### Determination of flavonoid in the roots

The total flavonoid content in roots was assessed using the spectrophotometric method (Piyanete et al. 2009). Briefly, 0.5 mL of a 70% ethanol extract derived from the roots was combined with 2 mL of distilled water and 0.15 mL of a 5% NaNO_2_ solution. This mixture was incubated for 6 min, after which 0.15 mL of a 10% AlCl_3_ solution was introduced, and then incubated for an additional 6 min. Subsequently, 2 mL of a 4% NaOH solution was added. The final volume was adjusted to 5 mL with methanol, and the mixture was mixed thoroughly. Following a 15 min incubation, the absorbance was recorded at 510 nm using a spectrophotometer. The total flavonoid concentration was calculated in milligrams per gram of dry weight, utilizing a quercetin standard curve for the determination.

### Amplicon sequencing of 16S and ITS microbial community analysis in the roots

Microbial populations in root samples were examined using Illumina amplicon sequencing, which targets the ITS region for fungus and the 16S rRNA gene for bacteria, where appropriate. The samples were vortexed in sterile phosphate-buffered saline (PBS) for 10 sec, followed by two rinses with sterile water. DNA was extracted from about 0.2 g of cleaned root tissue using the Wizard® Genomic DNA Purification Kit (Promega Corporation, USA). RNase and Proteinase K treatments were used during the extraction process to get rid of RNA and protein impurities, respectively. The primer pairs 341F (CCTACGGGNGGCWGCAG) and 805R (GACTACHVGGGTATCTAATCC) and ITS3 (GATGAAGAACGYAGYRAA) and ITS4 (TCCTCCGCTTATTGATATGC) were used to create the amplicon libraries for the 16S rRNA and ITS genes, respectively. Sequencing was performed on the Illumina Novaseq 6000 platform (PE250). Raw sequencing reads were processed and trimmed using Cutadapt (Martin, 2011), followed by analysis with the DADA2 pipeline (Callahan *et al.*, 2016). Reads associated with mitochondria or chloroplasts were excluded from the dataset. Taxonomic assignment of amplicon sequence variants (ASVs) was performed using the UNITE database (Nilsson *et al.*, 2019) for ITS regions and SILVA database version 138 (Quast *et al.*, 2013) for 16S rRNA sequences. Furthermore, microbial community composition was analyzed by calculating Bray-Curtis distance matrices from Hellinger-transformed data, followed by Principal Coordinate Analysis (PCoA) using the phyloseq and vegan packages in R. Amplicon sequence variants (ASVs) were classified at various taxonomic levels, and comprehensive analyses including alpha and beta diversity assessments, and relative abundance were performed using non-parametric Kruskal-Wallis tests determined at P < 0.05 (McMurdie and Holmes, 2013).

### Nanopore 16S bacterial community analysis in the nodules

Bacterial community analysis of the nodules was performed using 16S rRNA Nanopore sequencing. Briefly, root nodules were carefully exercised from soybean plants using sterile forceps and thoroughly washed under running tap water for 5 min to remove soil particles. The nodules were then immersed in 3% sodium hypochlorite for 3 min with occasional gentle shaking. The sterilized nodules were then rinsed three times with sterile distilled water before total DNA extraction was carried out using the CTAB procedure, as previously described (Clarke, 2009). For PCR experiments, 5 ng of DNA from each sample was combined with 16S primers 27F (AGAGTTTGATCMTGGCTCAG) and 1492R (CGGTTACCTTGTTACGACTT) to amplify the nearly full-length bacterial 16S rRNA gene. The PCR was performed under the following thermal cycling conditions: initial denaturation at 95°C for 2 min, followed by 30 cycles of denaturation at 95°C for 30 sec, annealing at 60°C for 30 sec, and extension at 72°C for 2 min. This was followed by a final extension at 72°C for 10 min and a hold at 4°C. Amplicons from each sample were ligated to pooled barcoded reads with the 16S Barcoding Kit 24 V14 (SQK-16S114-24, Oxford Nanopore Technologies, Oxford, UK) for library creation. The sequencing was done on a MinION portable sequencer (Oxford Nanopore Technologies) with a flow cell (FLO-FLG114). The MinION™ sequencing data (FASTQ files) were processed using the EPI2ME software from Oxford Nanopore Technologies, resulting in pass reads with an average quality score >9. The EPI2ME Fastq Barcoding method (Oxford Nanopore Technologies) was used to reduce adapter and barcode sequences. Principal coordinate analysis, alpha diversity, and relative abundance of the bacterial community were calculated using the R programs phyloseq and vegan. Non-parametric Kruskal-Wallis tests were used to detect variations in taxonomic abundance at a 5% significance level (McMurdie and Holmes, 2013).

### Determination of total soluble sugar in roots

The phenol-sulfuric acid method was employed as previously described with some modifications (Dey *et al.*, 1990). Briefly, 100 mg of fresh root tissue was extracted in 10 mL of ethanol and incubated at 60°C for 1 hour. The extract was transferred to a 25 mL volumetric flask, and the residue was re-extracted with ethanol. The combined extracts were diluted to 25 mL with ethanol. A 1 mL aliquot of the extract was mixed with 1 mL of 5% phenol solution in a thick-walled test tube, followed by the addition of 5 mL of concentrated sulfuric acid. The reaction mixture was thoroughly mixed and allowed to cool to room temperature. The absorbance of the resulting solution was measured at 485 nm using a spectrophotometer. Total soluble sugar concentration was quantified against a glucose standard curve and expressed as mg of glucose equivalents per gram of fresh weight (mg g ¹ FW).

### Determination of ammonium in roots

Ammonia concentrations in root tissues were determined using Nessler’s reagent spectrophotometric method (Jeong *et al.*, 2013). Approximately 100 mg of fresh root tissue was homogenized in 5 mL of 2% (w/v) potassium chloride solution, and the supernatant was collected after centrifugation at 12,000 rpm for 10 min. One milliliter of the extract was mixed with 1 mL of Nessler’s reagent and incubated at room temperature for 15 min. The absorbance of the reaction mixture was measured at 420 nm using a spectrophotometer. Ammonia content was quantified against a standard curve prepared with known concentrations of ammonium sulfate and expressed per gram of fresh weight (µg g ¹).

### Molecular detection of *V. paradoxus* and *P. lilacinus* in roots

The colonization efficiency of *V. paradoxus* and *P. lilacinus* in the roots was assessed using DNA-based qPCR. Briefly, root samples were washed twice with sterile phosphate-buffered saline, vortexed, and then rinsed twice with sterile water. DNA was extracted from approximately 0.2 g of root tissue using the cetyltrimethylammonium bromide (CTAB) method (Clarke, 2009). The extracted DNA was quantified using a NanoDrop ND-1000 Spectrophotometer (Wilmington, USA) and standardized by diluting all DNA samples to equal concentrations. The qPCR analysis was conducted using a CFX96 Touch Real-Time PCR Detection System (Bio-Rad, USA), with specific primers for *V. paradoxus* (forward: CAATCGTGGGGGATAACGC, reverse: GGCCGCTCCATTCGCGCA) and *P. lilacinus* (forward: CTC AGT TGC CTC GGC GGG AA, reverse: GTG CAA CTC AGA GAA GAA ATT CCG). The soybean β*-actin* (forward: GAGCTATGAATTGCCTGATGG, reverse: CGTITCATGAATTCCAGTAGC) that served as the internal control for relative quantification using 2-^ΔΔ^CT method (Livak and Schmittgen 2001).

### Microbial co-culture

We collected *P. lilacinus* (ATCC 10843) and *V. paradoxus* (B-1908) from the USDA Microbial Germplasm as representative strains of the genera *Paecilomyces* and *Variovorax*, respectively (Supplementary Fig. S3). We have tested the compatibility and interactions of *P. lilacinus* and *V. paradoxus* on nutrient agar in a series of *in vitro* time-course experiments. The microbial species (5 µl inoculum) were cultivated in three distinct combinations: *P. lilacinus* alone, *V. paradoxus* alone, and *P. lilacinus* and *V. paradoxus* for seven d. The growth of the microbial species was monitored for up to 7 d of culture using ImageJ software (National Institutes of Health, USA). The colony radius was measured at three different points and averaged to determine the growth of microbes.

### Split-root assay in Clark

Split-root experiments were conducted on autoclaved soil consisting of a 1:2 mixture of natural field soil and commercial potting mix. The plants were grown for 2 weeks before performing the split-root assay in soil pots. This assay was conducted using a single pot with a partitioned split-root assembly (compartment), with some modifications (Thilakarathna and Cope 2021). The taproot of each seedling was severed at the midpoint, and the lateral roots were evenly distributed between two compartments within the same pot, which were filled with a mixture of natural field soil and commercial potting mix in a 1:2 ratio. A solid plastic barrier was used to separate the compartments. Microbial inoculants (*V. paradoxus* and *P. lilacinus* at a concentration of 1×10 CFU per gram) were applied to one compartment of the split-root system, while the other compartment remained uninoculated, depending on the treatment (Fig. 8C). The treatments included three conditions: SR1: control/control (-/-), SR2: control/microbiome (-/+), and SR3: microbiome/microbiome (+/+). The split-root plants were then cultivated under these conditions for an additional four weeks before data collection.

### Data analysis

The statistical significance of each variable was evaluated using a combination of Student’s t-test for pairwise comparisons and analysis of variance (ANOVA) for evaluating differences across multiple groups, where appropriate. Where significant differences were detected by ANOVA, Duncan’s Multiple Range Test (DMRT) was employed as a post hoc analysis to identify specific group differences. All statistical analyses were conducted using SPSS Statistics 20.0 (IBM), ensuring a minimum of three biological replicates. Graphical representations were generated using the R package ggplot2.

## 3. Results

### Changes in morphological and physiological attributes

We observed distinct morphological and physiological responses in Clark and Arisoy under combined Fe deficiency and drought compared to untreated controls (Fig. 1A-1M). The combined stress caused a significant decrease in the number of nodules in both genotypes, with the decrease being more pronounced in Arisoy than in Clark (Fig. 1B). Both the root length and root dry weight were significantly declined due to combined Fe deficiency and drought in both genotypes relative to controls, with Clark showing a moderate reduction and Arisoy experiencing a more pronounced decline (Fig. 1C-1D). Shoot height and shoot dry weight showed no significant change in Clark but decreased significantly in Arisoy subjected to combined Fe deficiency and drought compared to controls (Fig. 1E-1F). Similarly, leaf relative water content (RWC) and SPAD score did not change significantly in Clark, whereas Arisoy showed a significant decrease in RWC and SPAD score under combined Fe deficiency and drought relative to controls (Fig. 1G-1H). Photosynthetic efficiency (Fv/Fm) was significantly impaired under Fe-drought+ conditions in both genotypes in contrast to untreated controls, with a less pronounced decline in Clark and a more severe reduction in Arisoy (Fig. 1I). In this study, H_2_O_2_ concentration showed no significant change in Clark but increased significantly in the roots of Arisoy under combined Fe deficiency and drought relative to controls (Fig. 1K). We found that Fe-drought+ did not significantly alter hydroxyl radical (•OH) levels in the roots. However, in Arisoy, Fe-drought stress led to a significant increase in •OH levels (*p* < 0.01) under concurrent Fe deficiency and drought relative to controls (Fig. 1K). The Fenton reaction rate showed contrasting responses in the two genotypes under Fe-drought+ conditions. In Clark, the Fenton reaction rate significantly decreased while Arisoy exhibited a significant increase in the Fenton reaction rate in the roots due to combined Fe deficiency and drought compared to controls (Fig. 1L). In Clark, ferric chelate reductase activity significantly increased, whereas, in Arisoy, it showed a marked decline in the roots under Fe-drought+ conditions compared to the controls (Fig. 1M).

**Fig. 1.**
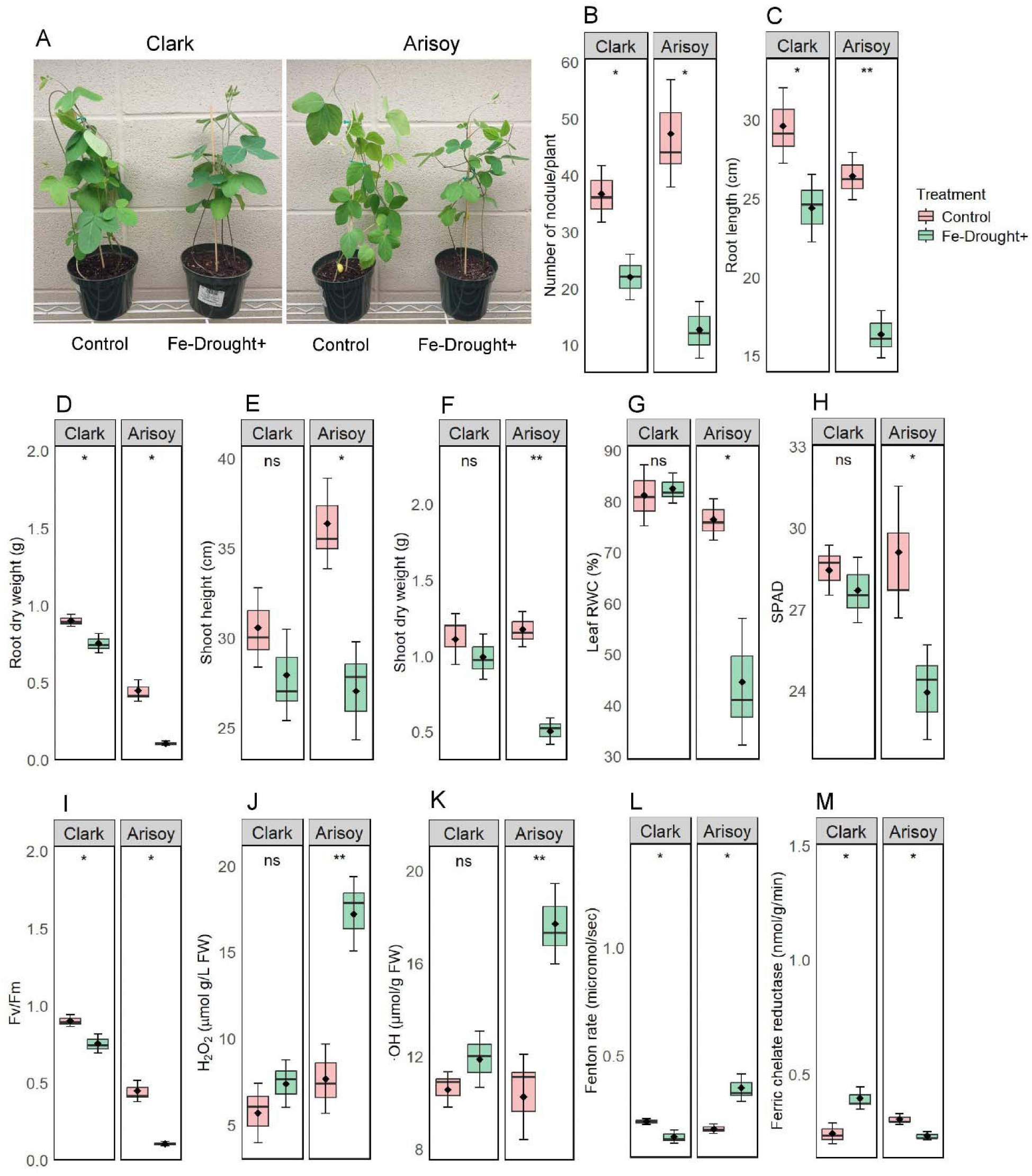
Growth, physiological, and biochemical responses of Clark and Arisoy genotypes under control and Fe-Drought+ conditions. A) aboveground phenotypes, B) number of nodules, C) root length, D) root dry weight, E) shoot height, F) shoot dry weight, G) leaf relative water content (RWC), H) SPAD chlorophyll content, I) photosynthetic efficiency (Fv/Fm), J) hydrogen peroxide (H_2_O_2_) in roots, K) hydroxyl radical (•OH) in roots, L) Fenton reaction rate in roots and M) ferric chelate reductase activity in roots. Data are presented as means ± SD (*n* = 3), with significance levels indicated by *\*P* < 0.05, \*\**P* < 0.01, \*\*\**P* < 0.001, and ns (not significant), based on a *t*-test.

We also cultivated a set of plants in sterile vermiculite conditions to study the effect of combined Fe deficiency and drought on growth parameters. Under Fe-Drought+ conditions, both Clark and Arisoy cultivars showed significant reductions in SPAD, shoot height, shoot fresh weight, shoot dry weight, and leaf RWC compared to the control (Supplementary Table S2). In Clark, stem diameter, number of leaves, and root length showed no significant differences between treatments. In Arisoy, all parameters, including stem diameter, number of leaves, and root length, showed significant differences between the control and Fe-drought+ conditions (Supplementary Table S2).

### Changes in nutritional status and symbiotic metabolites

Significant changes in nutrient content were observed in both roots and leaves of Clark and Arosy exposed to combined Fe deficiency and drought (Table 1). In roots, Fe, S and N contents remained unchanged, while Mn content significantly decreased subjected to Fe deficiency and drought relative to untreated controls. In Clark leaves, no significant changes were observed for Fe, Zn, or N due to stress while S and Mn showed a significant decrease (Table 1). In contrast, Arisoy exhibited more pronounced changes in both roots and leaves. In the roots, Fe, Mn, and N contents significantly decreased, while Zn and S showed no significant changes in response to combined Fe deficiency and drought. Furthermore, Fe, Mn, S, and N contents showed a significant decline in the leaves of Arisoy due to stress, but Zn levels remained unchanged (Table 1). Under stress, P content increased significantly in Arisoy roots, while no significant change was detected in Clark. In leaf, P levels remained unchanged in both genotypes under combined Fe deficiency and drought compared to controls (Table 1). We also found that flavonoid content significantly increased, while soluble sugar and ammonium contents showed no significant changes between control and Fe-drought+ conditions in the roots of Clark. On the flip side, Arisoy exhibited a significant decrease in flavonoid, soluble sugar, and ammonium contents under Fe-Drought+ conditions compared to the control (Table 2).

**Table 1.**
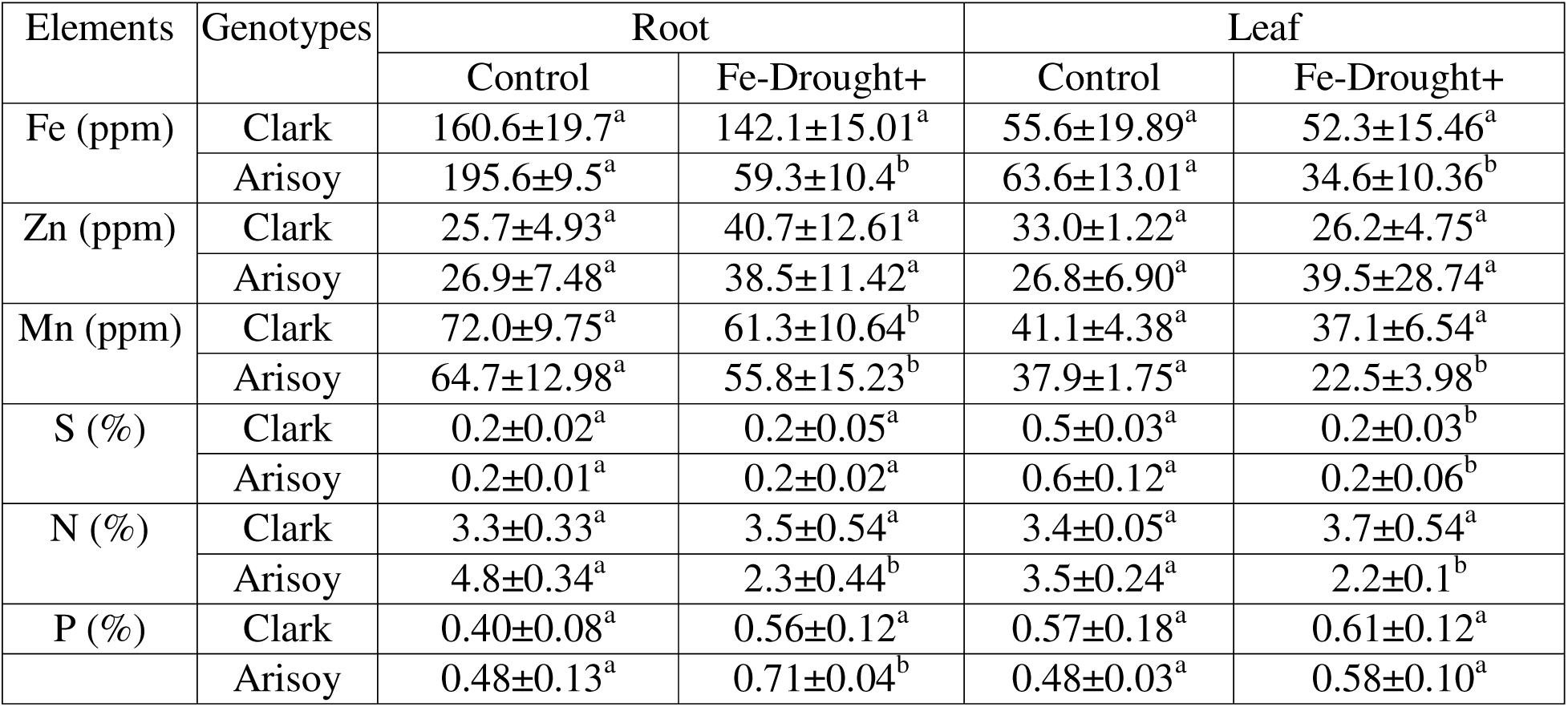
ICP-MS analysis of nutrient content in soybean roots and leaves under control and Fe-Drought+ conditions. Different letters denote significant differences between control and Fe-Drought+ conditions (*P* < 0.05, t-test). Data are presented as means ± SD from three independent biological samples (*n* = 3).

| Elements | Genotypes | Root |  | Leaf |  |
| --- | --- | --- | --- | --- | --- |
|  |  | Control | Fe-Drought+ | Control | Fe-Drought+ |
| Fe (ppm) | Clark | 160.6±19.7<sup>a</sup> | 142.1±15.01<sup>a</sup> | 55.6±19.89<sup>a</sup> | 52.3±15.46<sup>a</sup> |
| | Arisoy | 195.6±9.5<sup>a</sup> | 59.3±10.4<sup>b</sup> | 63.6±13.01<sup>a</sup> | 34.6±10.36<sup>b</sup> |
| Zn (ppm) | Clark | 25.7±4.93<sup>a</sup> | 40.7±12.61<sup>a</sup> | 33.0±1.22<sup>a</sup> | 26.2±4.75<sup>a</sup> |
| | Arisoy | 26.9±7.48<sup>a</sup> | 38.5±11.42<sup>a</sup> | 26.8±6.90<sup>a</sup> | 39.5±28.74<sup>a</sup> |
| Mn (ppm) | Clark | 72.0±9.75<sup>a</sup> | 61.3±10.64<sup>b</sup> | 41.1±4.38<sup>a</sup> | 37.1±6.54<sup>a</sup> |
| | Arisoy | 64.7±12.98<sup>a</sup> | 55.8±15.23<sup>b</sup> | 37.9±1.75<sup>a</sup> | 22.5±3.98<sup>b</sup> |
| S (%) | Clark | 0.2±0.02<sup>a</sup> | 0.2±0.05<sup>a</sup> | 0.5±0.03<sup>a</sup> | 0.2±0.03<sup>b</sup> |
| | Arisoy | 0.2±0.01<sup>a</sup> | 0.2±0.02<sup>a</sup> | 0.6±0.12<sup>a</sup> | 0.2±0.06<sup>b</sup> |
| N (%) | Clark | 3.3±0.33<sup>a</sup> | 3.5±0.54<sup>a</sup> | 3.4±0.05<sup>a</sup> | 3.7±0.54<sup>a</sup> |
| | Arisoy | 4.8±0.34<sup>a</sup> | 2.3±0.44<sup>b</sup> | 3.5±0.24<sup>a</sup> | 2.2±0.1<sup>b</sup> |
| P (%) | Clark | 0.40±0.08<sup>a</sup> | 0.56±0.12<sup>a</sup> | 0.57±0.18<sup>a</sup> | 0.61±0.12<sup>a</sup> |
|  | Arisoy | 0.48±0.13<sup>a</sup> | 0.71±0.04<sup>b</sup> | 0.48±0.03<sup>a</sup> | 0.58±0.10<sup>a</sup> |

**Table 2.** Changes in flavonoid, total soluble sugar and ammonium content in the roots of soybean grown in control and Fe-Drought+ conditions. Different letters indicate significant differences between the control and Fe-Drought+ conditions at a *P* <0.05 level, based on *t*-test. The data represents means ± SD of three independent biological samples (*n* = 3).

| Genotypes | Flavonoid (mg g<sup>-1</sup> DW) |  | Soluble sugar (mg g<sup>-1</sup> FW) |  | Ammonium (μg g<sup>-1</sup> FW) |  |
| --- | --- | --- | --- | --- | --- | --- |
|  | Control | Fe-Drought+ | Control | Fe-Drought+ | Control | Fe-Drought+ |
| Clark | 12.1±0.68<sup>a</sup> | 17.6±0.86<sup>b</sup> | 11.2±3.36<sup>a</sup> | 10.7±3.28<sup>a</sup> | 31.0±7.13<sup>a</sup> | 29.4±7.14<sup>a</sup> |
| Arisoy | 14.4±0.8<sup>a</sup> | 10.2±1.15<sup>b</sup> | 10.7±3.40<sup>a</sup> | 6.3±1.43<sup>b</sup> | 30.0±6.05<sup>a</sup> | 20.2±4.41<sup>b</sup> |

### Transcriptional changes in the roots

Principal component analysis (PCA) of transcriptomic data revealed distinct clustering between control and Fe-drought stress treatments for both Clark and Arisoy genotypes, indicating significant transcriptional reprogramming (Fig. 2A and 2B). In Clark, the first two principal components explained 57.5% and 27.2% of the variance, respectively, while in Arisoy, they accounted for 53.8% and 24.6% of the variance. Differential gene expression analysis identified a greater number of differentially expressed genes (DEGs) in Clark compared to Arisoy. Specifically, Clark exhibited 818 upregulated and 931 downregulated genes under Fe-drought+ conditions (Fig. 2C), while Arisoy showed 500 upregulated and 361 downregulated genes in the roots (Fig. 2D). The heatmap illustrates the differential expression of stress-responsive genes in two soybean genotypes, Clark and Arisoy, under combined Fe deficiency and drought stress compared to control conditions (Fig. 2E-G; Supplementary Table S4). In RNA-seq analysis, Clark exhibited higher expression levels of key genes associated with stress tolerance, such as *GLYMA.20G241500* (*Chalcone-flavone isomerase 1*), *GLYMA.08G059700* (*Sugar transporter 1*), *GLYMA.04G198400* (*Bidirectional sugar transporter SWEET10*), *GLYMA.05G035500* (*EamA-like transporter*) and *GLYMA.15G154100* (*2-oxoglutarate*) while Arisoy shows comparatively lower expression or downregulation of these genes in the roots in response to combined Fe deficiency and drought (Fig. 2E). The expression levels of genes related to nutrient uptake and transport including *GLYMA.10G146600* (Fe-regulated protein 3) and *GLYMA.10G167800* (*Ammonium transporter 1;2*), *GLYMA.07G006500* (*Sulfate transporter 3;4*) and *GLYMA.17G119400* (*Stabilizer of Fe transporter SufD*) were upregulated in Clark but showed no significant changes in Arisoy. However, *GLYMA.07G113500* (*ATPase 11*) was significantly upregulated in both Clark and Arisoy due to combined Fe deficiency and drought (Fig. 2F). Further, genes related to osmotic adjustment such as *GLYMA.17G027400* (Drought-induced-19 protein) showed significantly higher expression in both cultivars, with slightly elevated levels in Arisoy due to combined Fe deficiency and drought. In contrast, *GLYMA.11G106900* (Dehydration-associated protein) was significantly upregulated in Clark while Arisoy showed a non-significant higher expression in the root (Fig. 2G). In addition, the expression levels of genes associated with ROS detoxification such as *GLYMA.19G091400* (Ferritin-like family protein), *GLYMA.08G013800* (*Glutathione peroxidase 6*), *GLYMA.07G032600* (*Glutaredoxin 4*) and *GLYMA.13G172300* (*Peroxidase 19*) exhibited significantly higher expression in Clark while Arisoy showed no significant changes in the roots due to combined Fe deficiency and drought (Fig. 2H).

**Fig. 2.**
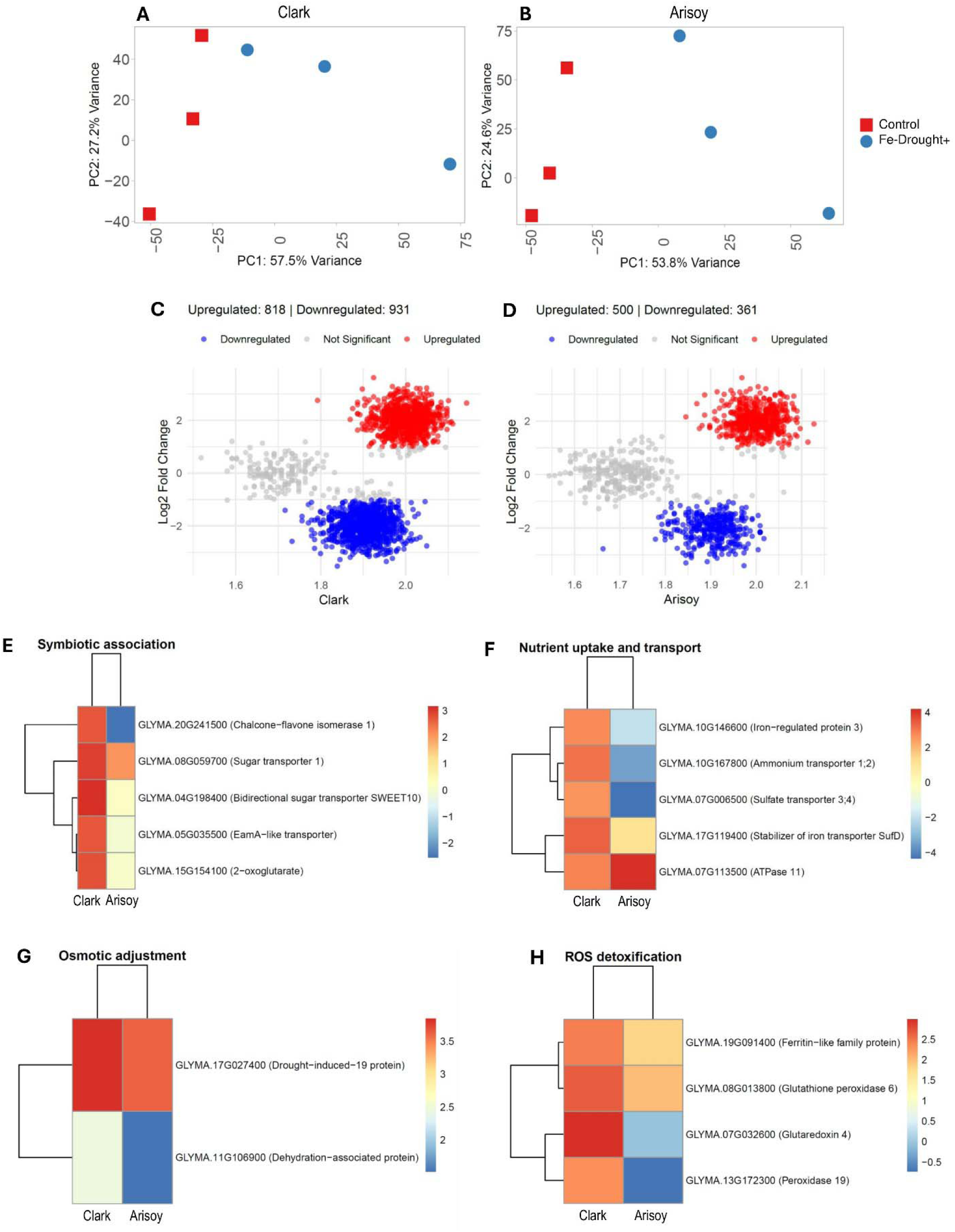
RNA-seq analysis in the roots of Clark and Arisoy under control and Fe-Drought+ conditions. (A, B) principal component analysis (PCA) in Clark (A) and Arisoy (B), volcano plots depicting differential gene expression in Clark (C) and Arisoy (D) with upregulated genes in red, downregulated genes in blue, and non-significant genes shown in gray and Heatmaps showing DEGs (log2 fold, P <0.05) related to key functional categories: symbiotic association (E), nutrient uptake and transport (F), osmotic adjustment (G), and ROS detoxification (H). Data represents three biological replicates, with color scales indicating the log2-fold changes in expression levels.

We further validated the RNA-seq results for a selected group of candidate genes using real-time PCR. The relative expression of *GLYMA.20G241500* (*Chalcone-flavone isomerase 1*), *GLYMA.04G198400* (*Bidirectional sugar transporter SWEET10*), *GLYMA.19G091400* (Ferritin-like family protein) and *GLYMA.10G146600* (Iron-regulated protein 3) was significantly increased in the roots of Clark under Fe-Drought+ conditions, while no significant difference was observed in Arisoy (Fig. 3A-3D). Furthermore, *GLYMA.17G027400* (Drought-induced-19 protein) showed a significant increase in both Clark and Arisoy exposed to Fe-Drought+ conditions compared to the control, while the increase was greater in Clark than in Arisoy (Fig. 3E).

**Fig. 3.**
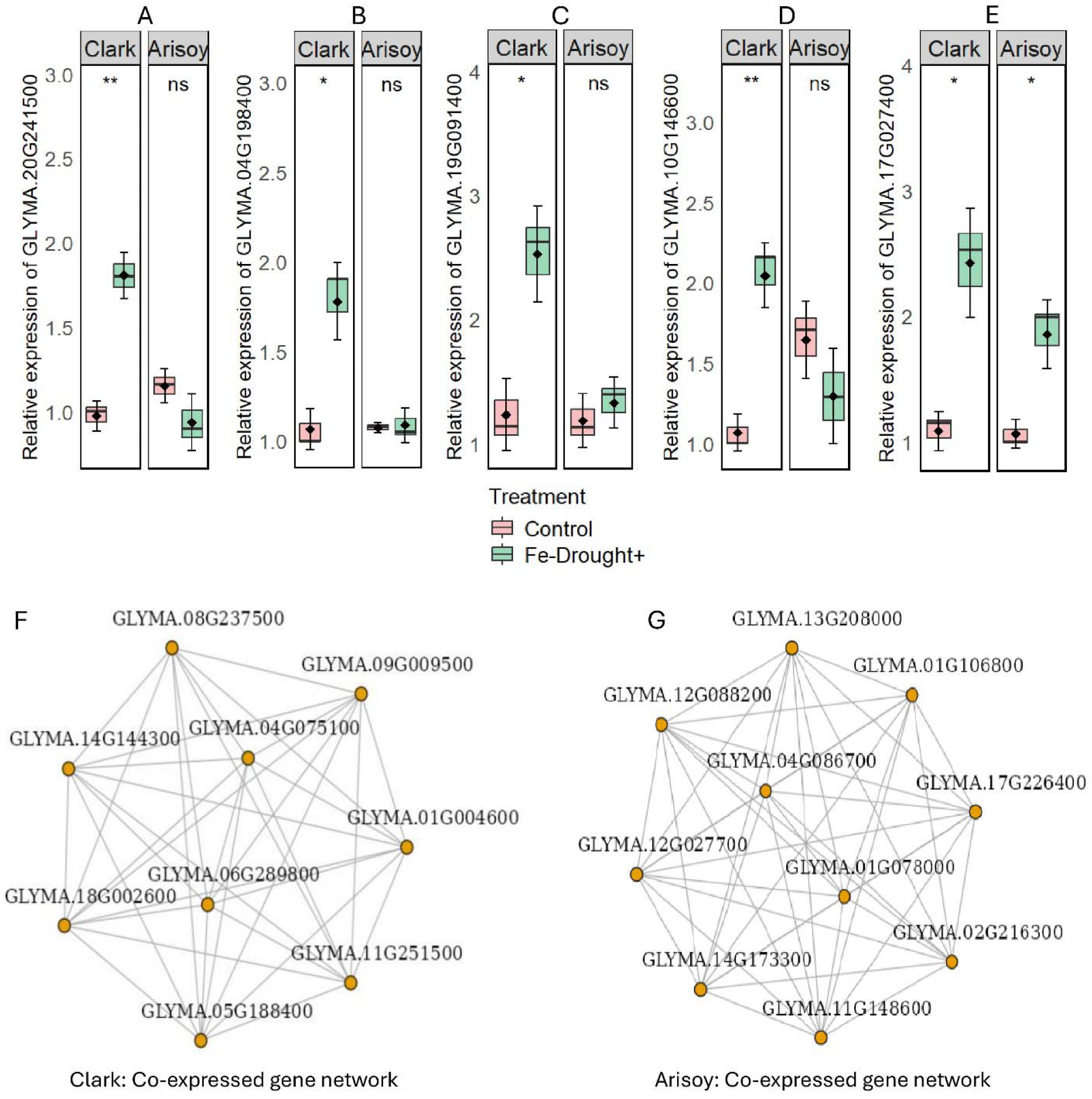
Expression of candidate genes (*GLYMA.20G241500*: *Chalcone-flavone isomerase 1*, *GLYMA.04G198400*: *Bidirectional sugar transporter SWEET10*, *GLYMA.19G091400*: Ferritin-like family protein, *GLYMA.10G146600*: Iron-regulated protein 3, *GLYMA.17G027400*: Drought-induced-19 protein) under controls and Fe-drought+ conditions and co-expressed gene network in the roots of Clark (*GLYMA.08G237500*: *DNA topoisomerase*, *GLYMA.09G009500*: *Cationic amino acid transporter*, *GLYMA.04G075100*: *Proton-translocating P-type ATPase*, *GLYMA.01G004600*: Unknown, *GLYMA.11G251500*: *Transcription factor SBP-box*, *GLYMA.05G188400*: Unknown, *GLYMA.06G289800*: *Pentatricopeptide repeat*, *GLYMA.18G002600*: *Exostosin-like*, *GLYMA.14G144300*: *Threonine-specific protein kinase*) and Arisoy (*GLYMA.13G208000*: *Aspartic peptidase*, *GLYMA.01G106800*: Small heat shock protein HSP20, *GLYMA.17G226400*: *Myosin*, *GLYMA.02G216300*: *Rhamnogalacturonan lyase*, *GLYMA.11G148600*: Unknown, *GLYMA.14G173300*: *Leucine-rich repeat*, *GLYMA.12G027700*: *Tetratricopeptide repeat*, *GLYMA.12G088200*: *Glycosyl transferase*, *GLYMA.04G086700*: *Threonine-specific protein kinase*, *GLYMA.01G078000*: Cysteine-rich repeat protein 15). Data are presented as means ± SD (*n* = 3), with significance levels indicated by *\*P* < 0.05, \*\**P* < 0.01, \*\*\**P* < 0.001, and ns (not significant), based on a *t*-test, where applicable.

The co-expression gene network analysis in the roots of Clark and Arisoy under Fe-Drought+ treatment revealed distinct transcriptional regulatory patterns. In Clark, key gene networks such as *GLYMA.09G009500* (*Cationic amino acid transporter*), *GLYMA.04G075100* (*Proton-translocating P-type ATPase*) and *GLYMA.14G144300* (*Threonine-specific protein kinase*) were highly interconnected, suggesting strong co-regulation in response to stress (Fig. 3F). In contrast, Arisoy showed the connection of other key gene networks such as *GLYMA.13G208000* (*Aspartic peptidase*), *GLYMA.01G106800* (Small heat shock protein HSP20) and *GLYMA.04G086700* (*Threonine-specific protein kinase*) (Fig. 3G).

The functional categorization and gene enrichment analysis showed distinct biological processes and KEGG pathways in Clark and Arisoy soybean genotypes under combined Fe deficiency and drought stress (Fig. 4A-3B). In Clark, the most enriched biological processes include detection of visible light, protein-chromophore linkage, polysaccharide catabolic process, response to water, carbohydrate metabolic process, oxidation-reduction process, and response to oxidative stress (Fig. 4A). For KEGG pathways, Clark shows significant enrichment in linoleic acid metabolism, alpha-linolenic acid metabolism, glycosphingolipid biosynthesis, circadian rhythm, starch and sucrose metabolism, and biosynthesis of secondary metabolites. In Arisoy, enriched biological processes include triglyceride biosynthetic process, detection of visible light, protein-chromophore linkage, glycerol ether metabolic process, polysaccharide catabolic process, and cell redox homeostasis. Furthermore, the enriched KEGG pathways in Arisoy include steroid biosynthesis, circadian rhythm, starch and sucrose metabolism, and glycerolipid metabolism (Fig. 4B).

**Fig. 4.**
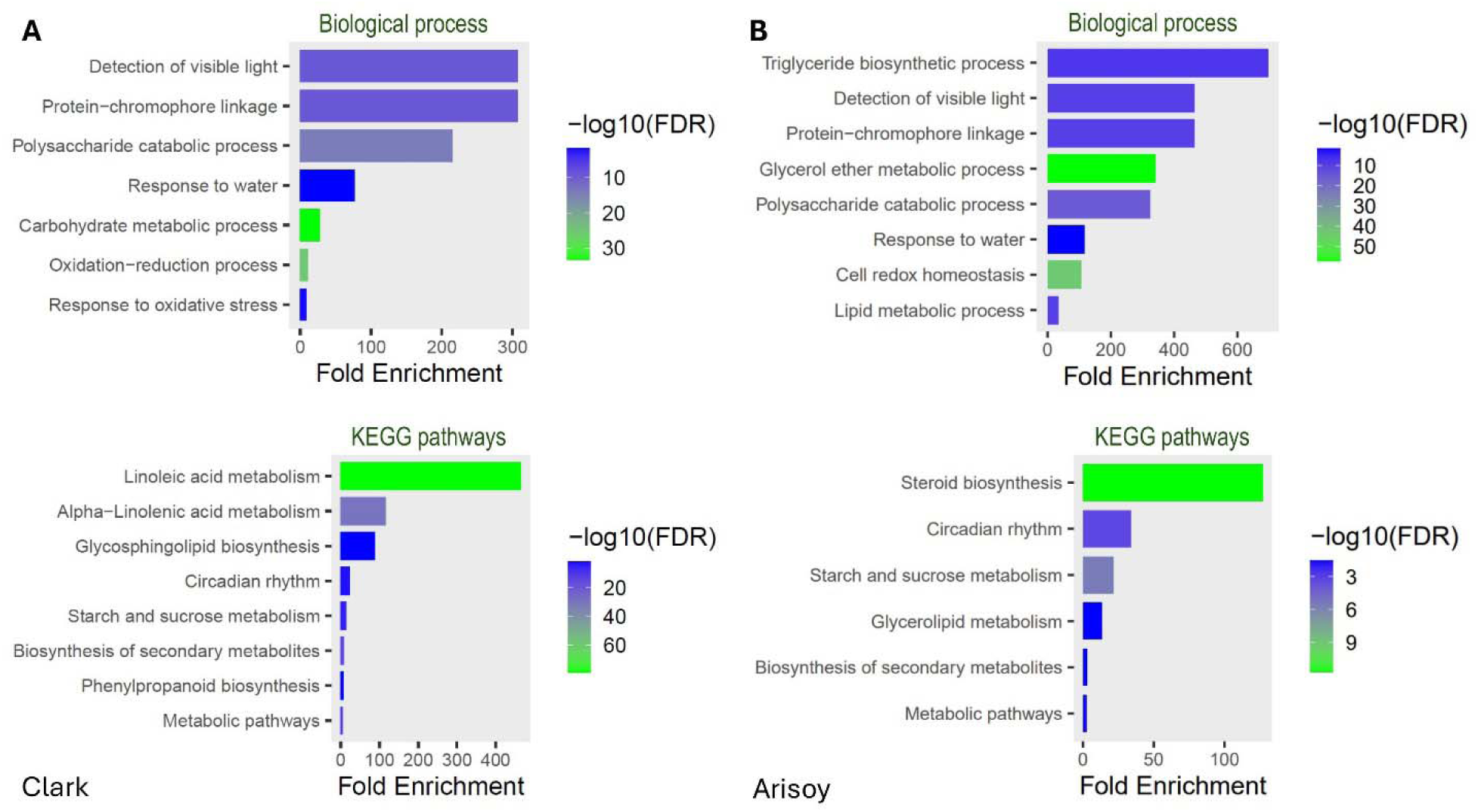
Functional enrichment analysis of biological processes and KEGG pathways in Clark and Arisoy under Fe-Drought+ stress. Enriched biological processes and KEGG pathways in Clark (A) and enriched biological processes and KEGG pathways in Arisoy (B). Data were analyzed using ShinyGO (V. 0.81), with fold enrichment and FDR significance shown on the plots.

### Changes in phytohormone levels in the roots

The phytohormonal profiles in the roots of Clark and Arisoy showed differential patterns in response to dual Fe deficiency and drought. Indole-3-acetic acid (IAA) levels significantly decreased under Fe-drought+ stress in both Clark and Arisoy due to Fe deficiency and drought relative to untreated controls (Fig. 5A). In contrast, abscisic acid (ABA) levels were significantly elevated in both genotypes under Fe-drought+ stress compared to controls (Fig. 5B). Jasmonic acid (JA) levels showed a significant increase in the roots of Clark but showed no significant changes in the Arisoy in response to combined Fe deficiency and drought compared to controls (Fig. 5C). The salicylic acid (SA) levels responded differently to Fe-drought+ in the contrasting soybean genotypes. In Clark, SA levels significantly increased under combined Fe deficiency and drought compared to controls. Conversely, in Arisoy, SA levels significantly decreased in such circumstances (Fig. 5D).

**Fig. 5.**
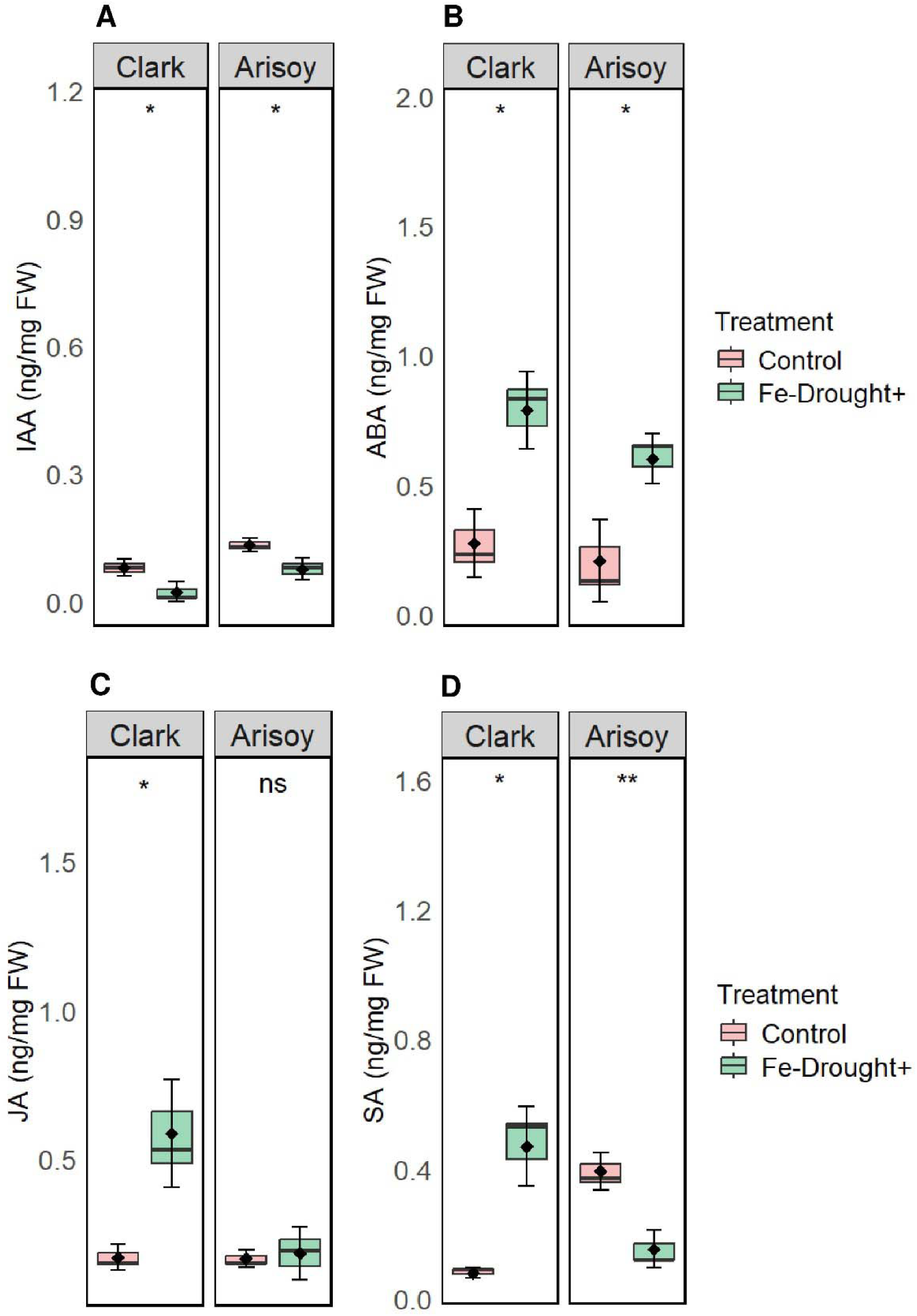
Hormonal profiles in the roots of Clark and Arisoy genotypes under control and Fe-Drought+ conditions. A) Indole-3-acetic acid (IAA), B) Abscisic acid (ABA), C) Jasmonic acid (JA), and D) Salicylic acid (SA). Data are presented as means ± SD of three biological replicates. Pink and green boxplots represent control and Fe-Drought+ conditions, respectively. Significant differences between treatments are indicated (*\*P* < 0.05, \*\**P* < 0.01; ns = not significant), based on a *t*-test.

### Shift in microbial dynamics in contrasting genotypes

We studied the effects of combined Fe deficiency and drought on the rhizosphere siderophore, microbial diversity, and community composition in the roots and nodules of Clark and Arisoy soybean genotypes. In Clark, Fe-drought+ stress significantly increased siderophore levels in the rhizosphere; however, no significant change was observed in Arisoy under the same conditions (Fig. 6A). The PCA analysis showed clear clustering of root 16S microbial communities between control and Fe-drought+ treatments in both genotypes (Fig. 6B). Alpha diversity metrics showed no significant changes in observed richness and Shannon diversity index in Clark under combined Fe deficiency and drought compared to controls. However, observed richness (*p* = 0.025) significantly decreased in Arisoy while Shannon diversity remained unaffected in Arisoy (Fig. 5C). The induction of Fe deficiency and drought in soil significantly altered the relative abundance of several genera in the root bacterial microbiomes. In Clark, the abundance of *Variovorax*, *Streptomyces*, and *Pararhizobium* increased significantly due to combined Fe deficiency and drought (Fig. 6D). In Arisoy, the abundance of *Streptomyces* and *Cellulomonas* significantly increased, while *Variovorax* and *Novosphingobium* significantly decreased in the roots in response to combined Fe deficiency and drought (Fig. 6E). The bacterial genera that showed no significant shift in relative abundance in the roots between control and Fe-drought+ conditions are as follows: Clark (*Shinella, Pseudomonas, Mycobacterium, Methylophilus, Herbaspirillum, Asticcacaulis*) and Arisoy (*Shinella, Pararhizobium, Mycobacterium, Methylophilus, Flavobacterium, Cellvibrio*). In this study, the nodule microbiomes were dominated by *Bradyrhizobium* in both genotypes under both conditions (Fig. 6F-6G). In Clark, *Bradyrhizobium* showed no significant changes, while in Arisoy, it significantly declined in relative abundance in the roots due to combined Fe deficiency and drought (Fig. 6F-6G). Furthermore, soybean root nodules from both the Clark and Arisoy cultivars exhibit distinct color differences under combined Fe deficiency and drought stress (Supplementary Fig. S2). In the control condition, the nodules displayed a vibrant pink color, whereas, under stress conditions, they showed a faded pink hue in both genotypes. However, the extent of color fading was less pronounced in Clark and more severe in Arisoy (Supplementary Fig. S2).

**Fig. 6.**
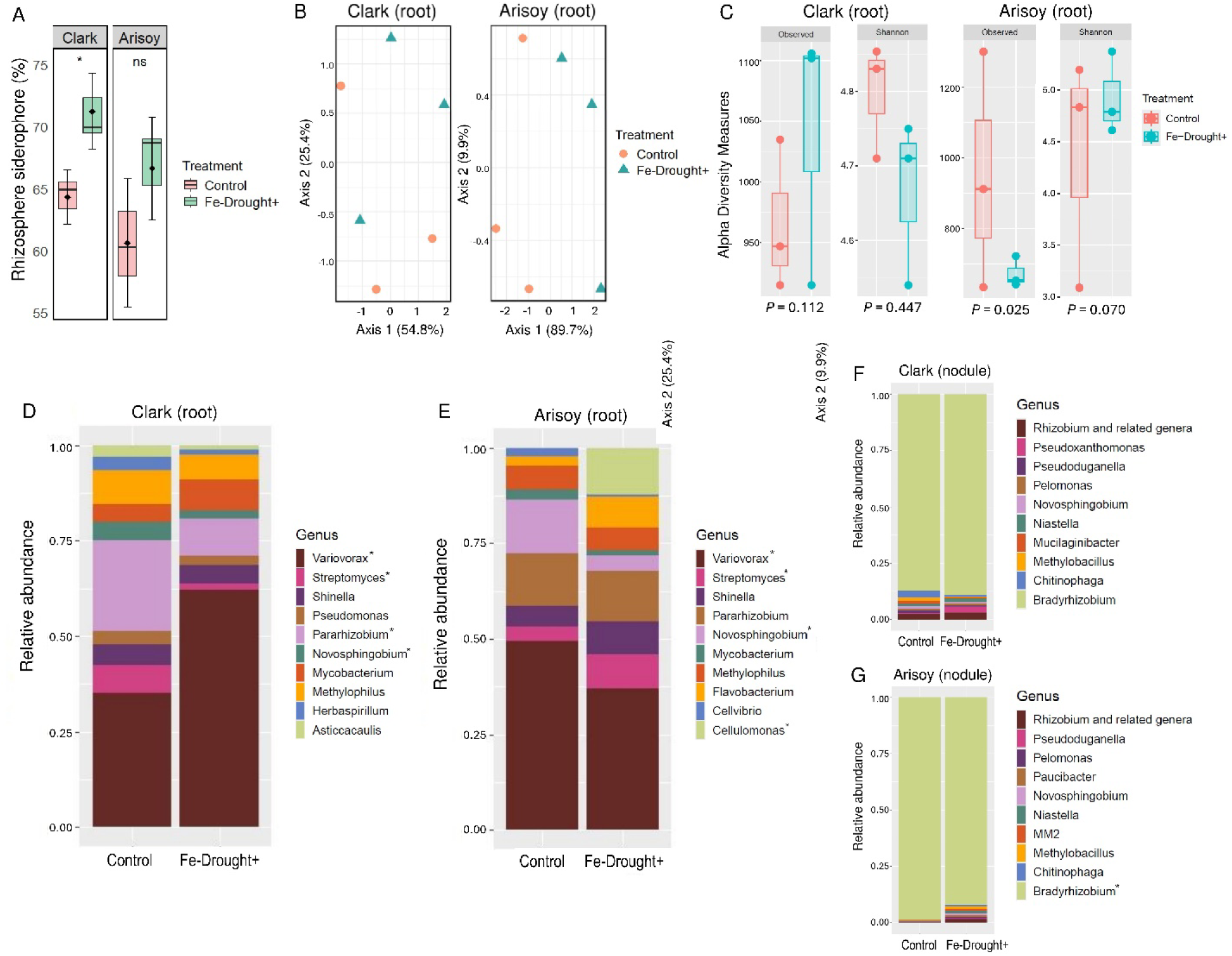
Siderophore levels and bacterial (16S) community composition and diversity in Clark and Arisoy under control and Fe-Drought+ conditions. Rhizosphere siderophore (A), principal component analysis (B), alpha diversity measures (C), the relative abundance of bacterial genera in the roots of Clark (D) and Arisoy (E), the relative abundance of bacterial genera in the nodules of Clark (F) and Arisoy (G). Asterisks (*) indicate significant differences based on the Kruskal-Wallis test (*P* < 0.05) based on three replicates (*n* = 3) per sample group.

In this study, the fungal community composition in the roots showed distinct clustering between the control and Fe-drought+ treatments in both genotypes, indicating shifts in fungal community composition under stress conditions (Fig. 7A). Alpha diversity measures (Panel B) showed no significant change in observed richness for either genotype under Fe-drought+ conditions (Fig. 7B). However, the Shannon diversity index significantly decreased in the roots of Clark under Fe-drought+ conditions, while no significant change was observed for Arisoy (Fig. 7B). Relative abundance analysis highlighted specific genera that were significantly impacted by Fe-drought+ conditions. In Clark roots, *Pseudohyphophila* and *Paecilomyces* significantly increased in abundance, while other genera, including *Trichocladium, Pseudoallescheria, Paracremonium, Humicola, Cladosporium, Chaetomium* and *Arnium,* showed no significant changes (Fig. 7C). In Arisoy roots, *Cladosporium* significantly increased, and *Paecilomyces* significantly decreased in abundance (Fig. 7D). Genera such as *Scedosporium, Pseudohyphophila, Pseudoallescheria, Podospora, Fusarium, Chaetomium, Ascospheara,* and *Arnium* remained unaffected by the Fe-drought+ treatment in the roots of Arisoy (Fig. 7D).

**Fig. 7.**
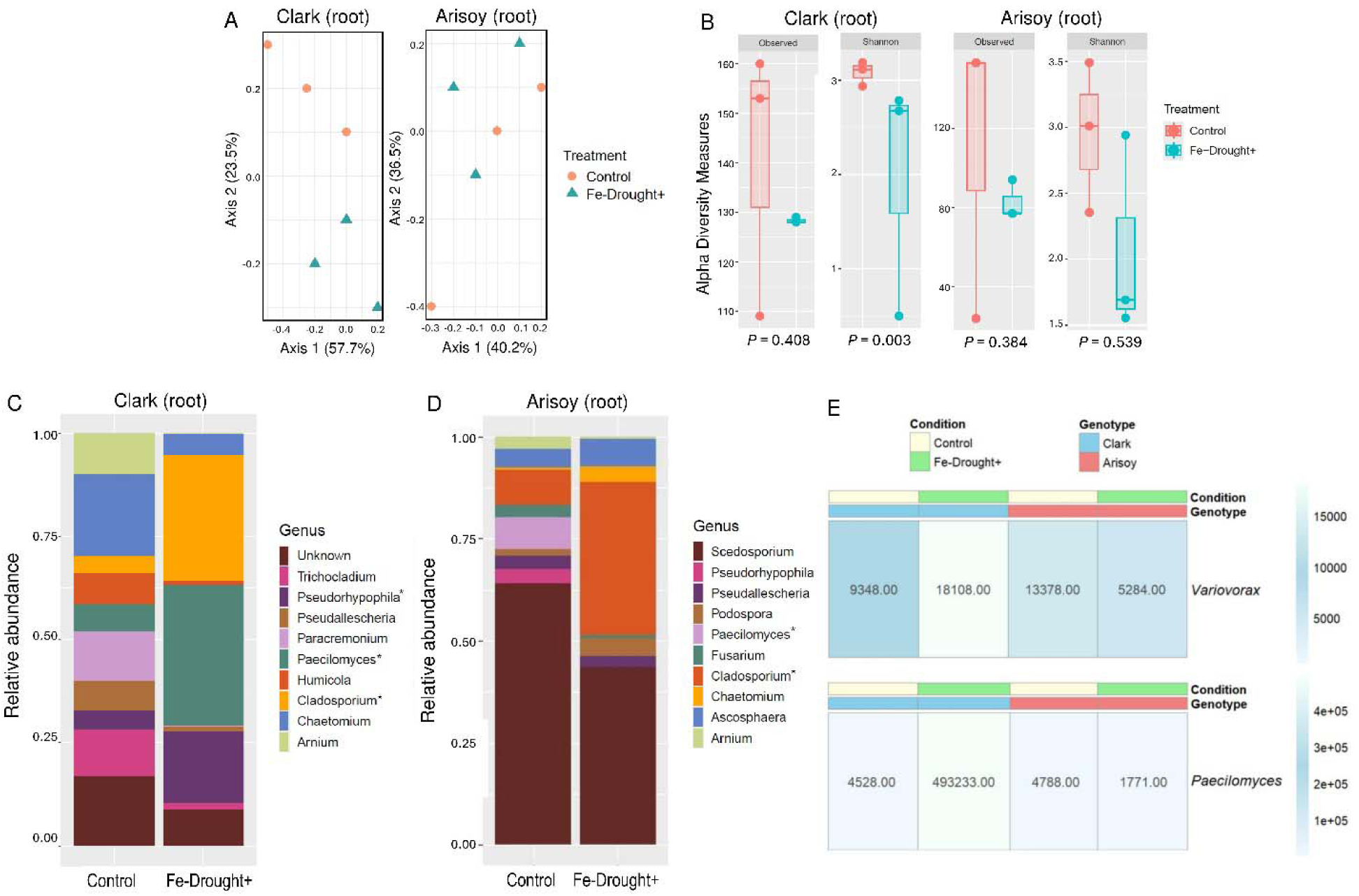
Fungal (ITS) community composition and diversity in the roots of Clark and Arisoy under control and Fe-Drought+ conditions. Principal component analysis (A), alpha diversity measures (B), the relative abundance of fungal genera in the roots of Clark (C) and Arisoy (D), and ASV abundance of *Variovorax* and *Paecilomyces* (E). Asterisks (*) indicate significant differences based on the Kruskal-Wallis test (*P* < 0.05) based on three replicates (*n* = 3) per sample group.

**Fig. 8.**
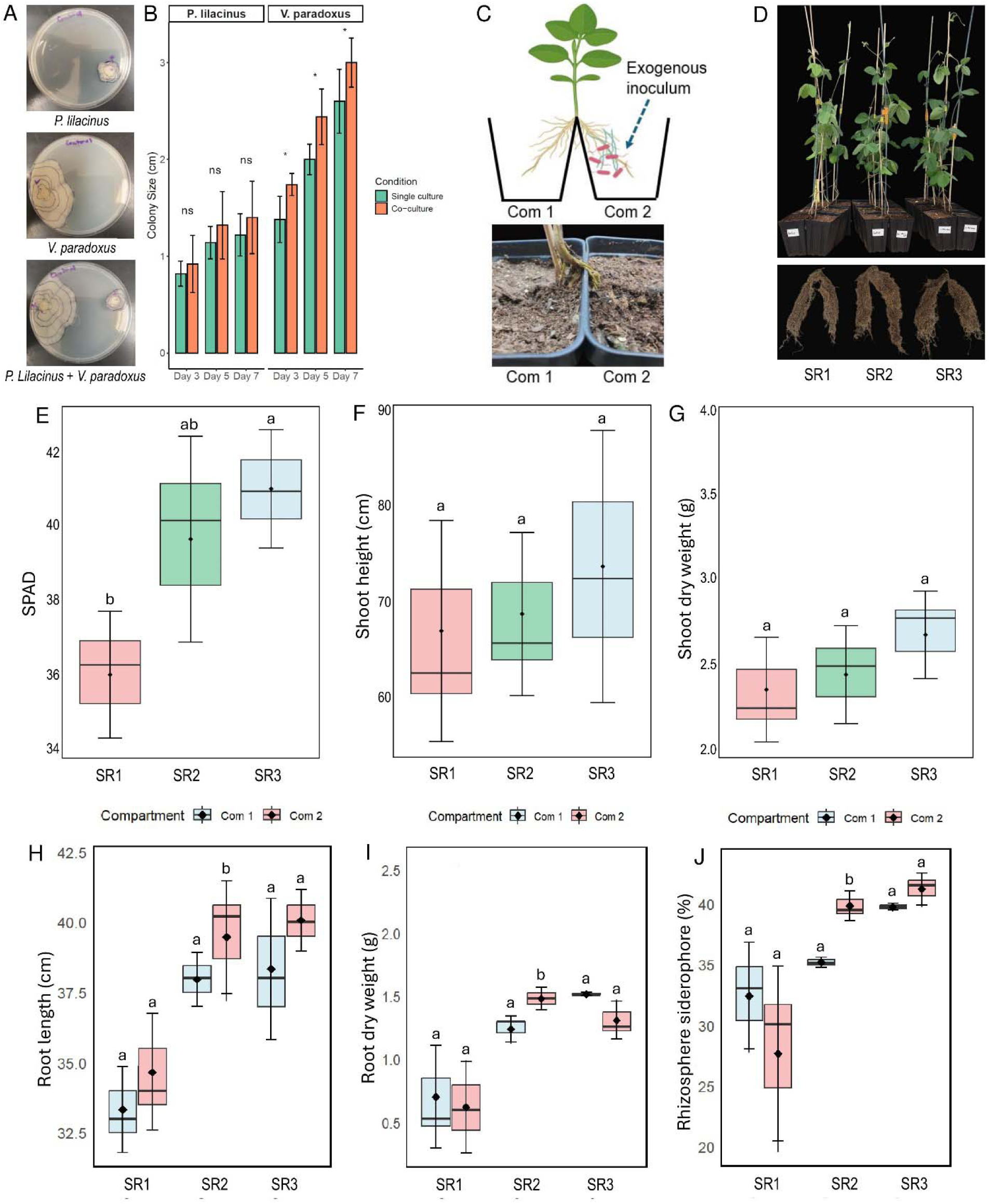
Interactions of enriched microbiomes and their effects on morpho-physiological responses in Clark under split-root systems. Colony growth of *Paecilomyces lilacinus* and *Variovorax paradoxus* in nutrient agar (A), colony size of *P. lilacinus* and *V. paradoxus* over time (B), diagram and establishment of split-root (SR) system (C), aboveground and belowground phenotype of split-root plants (D), split-root assay: SPAD chlorophyll content (E), shoot height (F), shoot dry weight (G), root length (H), root dry weight (I), and rhizosphere siderophore production (J). Asterisks (*) indicate significant differences (P < 0.05), and ’ns’ indicates non-significant differences (P > 0.05) in *Student’s t-test*. Furthermore, different letters indicate significant differences (*P* < 0.05, *n* = 3) between treatments based on ANOVA.

We further illustrated ASV counts for major beneficial microbes (*Variovorax* and *Paecilomyces*) across genotypes (Clark and Arisoy) differentially shifted due to combined Fe deficiency and drought (Fig. 7E). For *Variovorax*, the ASV number increased substantially in Clark under Fe-drought+ conditions (18,108 ASVs) compared to the control (9,348 ASVs). In contrast, Arisoy showed a substantial decrease in *Variovorax* between Fe-drought+ (5,284 ASVs) and control (13,378 ASVs) conditions. For *Paecilomyces*, the ASV count increased dramatically in Clark under Fe-drought+ conditions (493,233 ASVs) compared to the controls (4,528 ASVs). However, in Arisoy, *Paecilomyces* decreased under Fe-drought+ conditions (1,771 ASVs) compared to the control (4,788 ASVs), although the increase was less pronounced than in Clark (Fig. 7E).

### Interactions of *P. lilacinus* and *V. paradoxus* and changes in plant growth parameters in split-root plants

We found that there was no significant difference in colony size in *P. lilacinus* between single and co-culture conditions at all time points (Day 3, Day 5, and Day 7). In contrast, for *V. paradoxus*, the colony size in co-culture was significantly larger compared to single culture on all time points, indicating enhanced growth in the presence of *P. lilacinus* (Fig. 8A-8B). Furthermore, the split-root experiment was designed to evaluate the localized and systemic effects of enriched microbiomes on plant growth and physiology in Clark. Visual observations revealed that plants in the SR3 treatment (microbiome/microbiome) exhibited the most robust growth, with larger shoots and healthier root systems compared to SR1 (control/control) and SR2 (control/microbiome) systems (Fig. 8C-8D). In this study, SR1 exhibited significantly lower SPAD values compared to SR3, while SR2 showed intermediate values that were not significantly different from either SR1 or SR3 (Fig. 8E). However, shoot height and shoot dry weight remained stable across all treatments (Fig. 8F-8G). The root length and root dry weight showed significant differences between the compartments in SR2, while these root features showed no significant differences between the compartments in SR1 and SR3 (Fig. 8H-8I). Furthermore, the rhizosphere siderophore production showed significant differences between the compartments in SR2, with Com 2 having higher siderophore levels than Com 1. However, no significant differences were observed between the compartments in SR1 and SR3 (Fig. 8J).

### Effects of enriched microbiomes on sensitive soybean genotypes

The combined Fe deficiency and drought caused a significant decline in morphological and physiological traits of two sensitive genotypes (Arisoy and AG58XF3) of soybean (Fig. 9A-9F). We observed that inoculants consisting of enriched microbiomes (*V. paradoxus* and *P. lilacinus*) significantly increased leaf SPAD, plant height, plant biomass, leaf RWC, and root nodule number under combined stress compared to Fe-Drought+ conditions in both genotypes. Plants without stress but inoculated with the enriched microbiomes exhibited similar trends in these parameters to those of the controls in both Arisoy and AG58XF3 (Fig. 9A-9F). In Arisoy, rhizosphere siderophore showed no significant changes due to combined stresses but AG58XF3 showed a significant increase compared to controls (Fig. 9G). Interestingly, the addition of enriched microbiomes to the soil, whether in the presence or absence of combined stress, resulted in a significant increase in rhizosphere siderophore levels compared to Fe-Drought+ conditions. Siderophore levels were higher in the absence of stress than in its presence in both genotypes (Fig. 9G). Furthermore, flavonoid levels in the roots showed a significant decrease in both genotypes when plants were subjected to combined Fe deficiency and drought compared to untreated controls (Fig. 9H). However, flavonoid content in the roots of Arisoy showed a significant increase due to microbiome inoculation under stress, whereas no such increase was observed in AG58XF3 when compared to plants grown solely under stress. Both Arisoy and AG58XF3 inoculated with enriched microbiomes under control conditions exhibited flavonoid levels similar to those of the untreated controls (Fig. 9H).

**Fig. 9.**
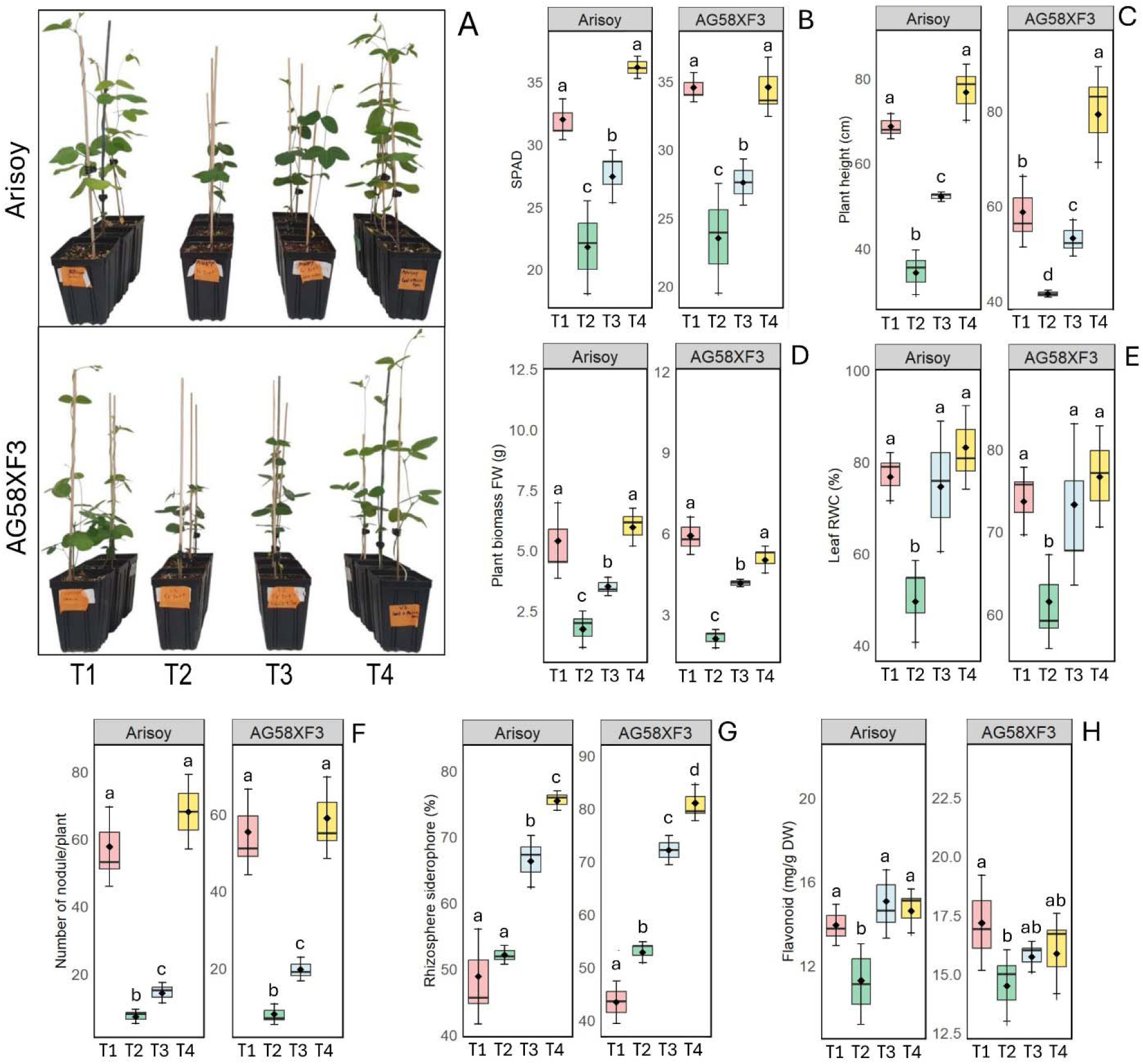
Effect of enriched microbiomes on soybean genotypes (Arisoy and AG58XF3): aboveground phenotype (A), leaf SPAD value (B), plant height (C), plant biomass (D), leaf relative water content (E), root nodule number (F), rhizosphere siderophore (G) and root flavonoid content (H). The treatments include T1: Control, T2: Fe-Drought+, T3: Fe-Drought+ Microbiome, and T4: Microbiome+. Different letters indicate significant differences among treatments within each genotype (*P* < 0.05). Data are presented as means ± SD from three independent replicates. The enriched microbiomes included *V. paradoxus* and P*. lilacinus*.

## 4. Discussion

Fe and water status are entwined in plants. Hence stress resilience in plants to the combined Fe deficiency and drought stress involves a complex interplay of physiological, molecular, and microbial associations (Santos *et al.*, 2023; Kanwar *et al.*, 2021). Our findings reveal the importance of tissue-level Fe retention, hormonal signaling pathways, and the enrichment of specific beneficial microbes in mitigating the adverse effects of dual stresses in soybean. This study uniquely integrates transcriptomic, phytohormonal, and microbial analyses to offer a comprehensive understanding of the mechanisms involved in improving soybean resilience under combined iron deficiency and drought stress.

### Regulation of Fe homeostasis and ROS scavenging underlying stress tolerance

This study reveals significant genotype-specific variations in stress resilience between Clark and Arisoy under combined Fe deficiency and drought. Clark exhibited moderate reductions in root length and biomass while maintaining shoot biomass and leaf RWC under stress. Variations in RWC are closely linked to the genotypic background, which affects the ability of plants to tolerate drought stress (Zhang *et al.*, 2023; Kabir *et al.*, 2015). Therefore, the physiological adjustments, such as sustained water retention supported by stable photosynthetic efficiency, might have been effective in mitigating the impact of dual stress in Clark. This aligns with the previous findings linking chlorophyll stability and photosynthetic efficiency to withstand abiotic stress (Therby-Vale *et al.*, 2022; Wang *et al.*, 2018). The physiological adjustments of Clark are supported by transcriptomic data, which reveal the upregulation of genes associated with osmotic adjustment. For instance, the enhanced expression of drought-induced-19 protein and dehydration-associated protein in Clark may play a critical role in maintaining cellular integrity under drought. The overexpression of drought-induced genes improved drought and salt tolerance in various plants (Wu *et al.*, 2022; Wu *et al.*, 2020). These proteins also play a role in ABA-dependent signaling pathways and regulate stomatal aperture, The Fe homeostasis in stress resilience has been well-documented, thereby influencing plant water status during drought conditions (Wu *et al.*, 2020). In another study, DREB (dehydration-responsive element-binding protein) mutants showed increased sensitivity to drought in rice (Wang *et al.*, 2022).

A lack of water or minerals in soil often triggers the imbalance of nutrient status and overproduction of ROS in plants (Barzana *et al.*, 2021; Jiang and Zhang, 2002). In this study, the upregulation of the genes related to dehydration protection in Clark is consistent with the maintenance of mineral status, such as Fe and other nutrients, under combined stresses. The improved Fe status in Clark was further supported by the induction of Fe-regulated protein 3 in the roots. Fe regulatory proteins (IRPs) maintain Fe homeostasis in eukaryotic cells by binding to Fe-responsive elements in target mRNAs and regulating their stability (Zhang *et al.*, 2014). This upregulation of Fe-regulated protein 3 suggests a dual role not only as a Fe-storage protein but also as an active antioxidant defense component under combinatorial stress conditions. Fe-regulatory proteins, including both positive and negative regulators, work in coordination to maintain Fe homeostasis in plants exposed to Fe deficiency (Lichtblau et al. 2022). Furthermore, root Fe re-utilization and transport from root to shoot have been observed in Arabidopsis exposed to drought (Lei *et al.*, 2014). Thus, Fe-regulated proteins may serve as valuable markers of stress tolerance and potential targets for soybean improvement through breeding programs.

Fe availability is also crucial for enzymatic functions and photosynthetic machinery (Connorton *et al.*, 2017; Kobayashi and Nishizawa, 2012). Furthermore, elevated ATPase and FCR activities under Fe deficiency are closely linked to enhanced Fe acquisition, forming a key component of the Strategy I response in plants (Kabir *et al.*, 2012; Santi and Schmidt, 2009). In this study, the upregulation of *ATPase 11*, responsible for proton extrusion (H^+^), in both Clark and Arisoy suggests an adaptive response to promote rhizosphere acidification for mobilizing soluble Fe and other minerals. Interestingly, we also found a significant upregulation of *Sulfate transporter 3;4* and *Stabilizer of Fe transporter SufD* in Clark when cultivated in dual Fe deficiency and drought.

SufD is a key component of the SUF (S utilization factor) machinery essential for Fe-S cluster assembly and Fe acquisition in plants (Bai *et al.*, 2018). S plays a key role in maintaining the homeostasis of micronutrients such as Fe, Cu, Zn, and Mn primarily through S-derived organic ligands that facilitate metal uptake and translocation (Chorianopoulou and Bouranis 2022). Furthermore, Fe deficiency increases the demand for S uptake and assimilation (Astolfi *et al.*, 2021). These findings align with improved S status in the root, suggesting that retaining Fe and other micronutrients helps soybean plants cope with combined Fe deficiency and drought. In this study, the increase in phosphorus levels in the roots of Arisoy but not in Clark is surprising. One possible explanation is that in Fe-deficient soils, the reduced availability of soluble Fe³ limits the formation of FePO, a poorly soluble compound (Saga *et al.*, 2018. As a result, more P may remain available in the soil, leading to increased uptake. This observation was not observed in Clark, likely because several strategies related to Fe solubilization (e.g., *ATPase 11* expression and ferric chelate reductase activity) were induced in response to the combined stress of Fe deficiency and drought.

Furthermore, the ability of Clark to limit stress-induced damage was supported by reduced Fenton reactions in the roots. This aligns with the upregulation of Ferritin-like family protein in the roots of Clark exposed to simultaneous Fe deficiency and drought. Fe and ferritin are vital for drought tolerance as they support antioxidant enzymes in neutralizing ROS during stress (Zhang *et al.*, 2023; Ndou *et al.*, 2023). Ferritin, a Fe-storage protein, also acts as a central player to prevent the excessive accumulation of free Fe that could lead to ROS production (Briat *et al.*, 2010). This aligns with our findings, as the lower ROS such as H_2_O_2_ and •OH levels in Clark further support enhanced capacity in Clark to mitigate oxidative stress in concurrent stresses. In contrast, dual stress imposed a greater oxidative burden on Arisoy, possibly exacerbated by impaired water transport and stress signaling pathways. Additionally, differences in oxidative stress responses, particularly those associated with the Fenton reaction, may contribute to the contrasting resilience observed in Clark and Arisoy. The Fenton reaction is a chemical process in which Fe(II) reacts with H_2_O_2_ to generate highly reactive •OH in plants (Smirnoff and Arnaud, 2019; Richards *et al.*, 2015). This was likely due to its enhanced FCR activity and effective Fe utilization, which limited the availability of free Fe for ROS generation in Clar under stress. Reduced Fenton activity in Clark not only minimized oxidative damage but also allowed for better allocation of Fe to essential metabolic processes. These findings align with the role of Fe homeostasis and ROS scavenging in plant stress tolerance (Taheri *et al.*, 2022). The reductions in H_2_O_2_ and •OH levels were correlated with the upregulation of several ROS-related genes such as *Peroxidase 19*, *Glutathione peroxidase* and *Glutaredoxin 4* in Clark. Peroxidase genes mainly scavenge H_2_O_2_ resulting in the inhibition of •OH generations through the Fenton reaction in plants under abiotic stresses (Zhang *et al.*, 2022; Kidwai *et al.*, 2020). Transgenic plants expressing glutathione peroxidase genes (GPX) exhibited increased tolerance to drought, and oxidative damage in several plant species (Hou *et al.*, 2024; Zhang *et al.*, 2019). Furthermore, glutaredoxin protein plays a vital role in the assembly and maintenance of Fe-S clusters, which are essential for ROS detoxification and various metabolic processes (Couturier *et al.*, 2015). *Glutaredoxin* knockouts in plants have been shown to impair stress tolerance and increase oxidative damage (Laporte *et al.*, 2012). As a result, the better adaptability of Clark to Fe deficiency and drought is at least partially attributed to the expression of antioxidant and ROS-scavenging mechanisms, which help limit stress-induced cellular damage.

### Changes in hormonal balance and symbiotic determinants in the roots

Significant hormonal changes were observed in Clark and Arisoy under the conditions of combined Fe deficiency and drought. Both genotypes showed reduced IAA, probably reallocating resources from growth to stress adaptation. A decline in auxin levels under drought and nutrient deficiency showed the conservation of energy and resources for survival rather than growth (Kurepa and Smalle, 2022; Peleg and Blumwald, 2011). Interestingly, IAA reduction is often linked to a shift in the balance of phytohormones, favoring ABA and ethylene pathways in plants exposed to drought (Mubarik *et al.*, 2021; Gupta *et al.*, 2017). In this study, the induction of ABA in both genotypes suggests a general adaptive response of plants to regulate stomatal closure and limit water loss under drought (Munemasa *et al.*, 2015). Interestingly, JA levels were significantly elevated in Clark under Fe deficiency and drought, while Arisoy exhibited no changes. The interplay between ABA and JA is not conclusive and depends on the type of stress and plant species. However, JA accumulation is essential for ABA accumulation following drought treatment in *Arabidopsis* (de Ollas *et al.*, 2015). Similarly, increased JA levels promote ABA production in rice under drought stress (Kim *et al.*, 2009; Li *et al.*, 2019) suggesting that JA acts upstream of ABA in the regulatory pathway. Hence, the elevation of JA levels suggests that it may be one of the drivers of stress responses in Clark under combined Fe deficiency and drought. Additionally, Clark showed an increase in SA levels, which is known to trigger antioxidant genes and reduce ROS accumulation during drought stress in plants (Jahan *et al.*, 2023). These findings emphasize the role of hormonal interactions, particularly JA and SA induction in Clark, in enhancing tolerance to combined Fe deficiency and drought.

In our RNA-seq analysis, genes associated with symbiotic interactions, including *Chalcone-flavonone isomerase*, *2-Oxoglutarate*, *Sugar transporter protein 1*, and *Bidirectional sugar transporter SWEET10*, showed higher expression levels in the Clark genotype compared to Arisoy. Flavonoids, including chalcones and flavones, are key host determinants that regulate microbial recruitment and gene expression, thereby promoting symbiotic interactions (Tohge *et al.*, 2017; Kang *et al.*, 2014; Hassan and Mathesius, 2012). In our dataset, *Chalcone-flavone isomerase 1*, and *Sugar transporter 1* was upregulated in Clark but not in Arisoy under combined stress. These genes did not appear as strongly induced in Fe-only or drought-only, suggesting a possible synergistic response unique to dual stress conditions. However, the induction of chalcone-flavone isomerase enzyme is associated with anthocyanin biosynthesis, which has been reported to be triggered by Fe deficiency in grapevine (Caramanico *et al.*, 2017). Additionally, sucrose and hexoses released into the rhizosphere serve as energy sources, signaling molecules to attract beneficial microbes (Kryukov *et al.*, 2021). Particularly, Sugar transporter protein 1 and *Sugar efflux transporter SWEET10* direct carbon flow to the rhizosphere (Chen *et al.*, 2012) and support the energy demands of nitrogen-fixing symbionts (Breia *et al.*, 2021; Kryukov *et al.*, 2021). Also, the upregulation of the *EamA-like transporter gene* suggests the stability of Clark in nitrogen assimilation, as this transporter is known to contribute to nutrient exchange and signaling during symbiotic interactions in legumes (Garcia *et al.*, 2023; Banasiak *et al.*, 2021; Guefrachi *et al.*, 2015). Taken together, our findings suggest the strong connection of symbiotic benefits as a key factor in improving resilience to combined Fe deficiency and drought stress in soybean. The genotype-specific responses observed in Clark and Arisoy suggest that similar genes and microbial interactions could be explored in other legume crops.

### Shift in microbial recruitment underlying stress resilience

To further elucidate microbial associations with differential stress resilience, we performed 16S and ITS sequencing on roots and nodules. The microbial community dynamics in the nodule and roots also demonstrated genotype-specific dynamics in the microbiome. Microbial diversity acts as a crucial mediator between plants and their surrounding environment (Zhang *et al.*, 2021). In this study, the significant increase in the rhizosphere siderophore in Clark under Fe deficiency and drought suggests enhanced microbial activity aimed at improving Fe solubility and availability. Siderophore production is a known adaptive response of rhizosphere microbes to Fe-limiting environments (Kabir and Bennetzen, 2024). Hence, the lack of a similar response in Arisoy may indicate either limited microbial activity or insufficient recruitment of siderophore-producing microbes, which could partially explain its reduced tolerance to combined stress. This is supported by the stability of microbial abundance and diversity in Clark under stress. However, Arisoy exhibited a significant decline in bacterial abundance, indicating reduced microbial adaptability. This discrepancy in the bacterial community might contribute to the observed differences in stress tolerance between the genotypes. Furthermore, microbial diversity is linked to plant fitness, nutrient cycling, and stress mitigation (Naylor *et al.*, 2017).

Our study demonstrates how contrasting genotypes recruit and sustain beneficial microbes in response to Fe deficiency and drought. Several bacterial genera such as *Variovorax*, *Streptomyces*, *Pararhizobium*, and *Novosphingobium* were altered in the roots of Clark. The increase in *Variovorax* in Clark is particularly promising, given its well-documented role in nutrient solubilization and phytohormone modulation (Jiang *et al.*, 2012; Sun *et al.*, 2018). Also, the differential shift in *Streptomyces* between contrasting genotypes suggests a potential genotype-specific recruitment or interaction mechanism to stress. Previous studies reveal that drought-induced Fe limitation in sorghum roots disrupts Fe homeostasis, enriching *Streptomyces* in the rhizosphere (Xu *et al.*, 2021). They also found that the addition of exogenous Fe disrupts this enrichment and diminishes the plant growth-promoting effects of *Streptomyces*. Additionally, drought-responsive plant metabolism improves Monoderm bacterial lineages, including *Actinobacteria*, *Chloroflexi*, and *Firmicutes* (Xu and Coleman-Derr, 2019). The disappearance of *Streptomyces* in the roots under combined Fe deficiency and drought may be associated with the improved Fe status in Clark, aligning with findings in drought-exposed sorghum (Xu *et al.*, 2021). Furthermore, Clark may possess a genotype-specific ability to maintain Fe homeostasis, reducing the need for *Streptomyces* enrichment while the stress conditions could exacerbate Fe scarcity, intensifying the recruitment of *Streptomyces* in Arisoy.

As expected, *Bradyrhizobium* dominated the nodule microbiomes of both soybean genotypes. However, its significant decline in Arisoy under stress indicates disrupted symbiotic efficiency. This could result in reduced N availability, reflected in the N status of roots and shoots, whereas Clark exhibited no such changes. Particularly, drought directly affects key processes involved in nodulation by reducing soil moisture, which limits the mobility of nitrogen-fixing bacteria and impairs their ability to colonize legume roots (Lumactud *et al.*, 2023; Larrainzar *et al.*, 2007). Clark and Arisoy exhibited differences in the composition of minor genera, such as *Pseudoxanthomonas* and *Mucilaginibacter*. These differences are likely shaped by the host plant’s genetic makeup in response to stress conditions. Although Clark exhibited a reduction in nodule number, the abundance of *Bradyrhizobium* within the nodules remained stable and active, indicated by the pink-red color of the cross-sections under combined Fe deficiency and drought. The pink-red interior color of root nodules due to the presence of leghemoglobin may be associated with high rates of nitrogen fixation (Jiang *et al.*, 2021; Küçük and Cevheri, 2014). Moreover, the efficiency of Clark to maintain *Bradyrhizobium* supports its symbiotic stability, aligning with ammonium retention coupled with the upregulation of *Ammonium transporter 1* and N status. Similar studies have shown that Rhizobia-legume symbiosis is crucial for mitigating drought stress and enhancing nitrogen fixation in *Vicia sativa* and *Pisum sativum* (Álvarez-Aragón *et al.*, 2023). Hence, the stress resilience in Clark probably stems from active *Bradyrhizobium*, enabling ammonium production, which is subsequently imported into the roots via *Ammonium transporter 1*. In ITS analysis, genotypic differences in fungal communities in the root were evident with *Paecilomyces* significantly enriching in Clark but not in Arisoy. *P. variotii* functions as an elicitor, stimulating the growth of *Arabidopsis thaliana* by inducing auxin accumulation and improving disease resistance through the activation of the SA signaling pathway (Lu *et al.*, 2019). Similarly, tomato plants inoculated with *P. lilacinus* have been shown to exhibit enhanced growth parameters and increased flavonoid content (Musa *et al.*, 2023). Taken together, the differential microbial dynamics observed in Clark and Arisoy underline the critical role of the root-associated microbiomes in differential stress tolerance to Fe deficiency and drought. The ability of Clark to recruit *Variovorax* and *Paecilomyces* may pose the potential of leveraging enriched microbiomes as a key trait for selection to improve tolerance in soybean in response to combined Fe deficiency and drought.

### Interactions of enriched microbiomes, signaling pathways, and effects on inducing stress resilience

Microbial compatibility is essential for the success of microbial consortia in promoting plant growth (Prigigallo *et al.*, 2023). Microbial co-cultures can significantly alter the growth and metabolism often mediated by diverse metabolites produced by participating organisms (Bertrand *et al.*, 2014; Wolter and Hoffman, 2014). In this study, the increased colony size of *V. paradoxus* in the presence of *P. lilacinus,* suggests a beneficial effect of *P. lilacinus* on *V. paradoxus*. Such interactions are commonly observed in microbial communities, where one species facilitates the growth of another through metabolic complementation (Liu *et al.*, 2024; Mendes *et al.*, 2013). Furthermore, the lack of significant variation in the colony size of *P. lilacinus* across co-culture conditions suggests that *P. lilacinus* is either unaffected by or tolerant of the presence of *V. paradoxus*, maintaining its growth independently of any potential microbial interactions. The interactions between *V. paradoxus* and *P. lilacinus* with other microbes need further investigation. However, positive growth-promoting interactions are common among microbes, particularly between strains with different carbon consumption profiles (Kehe *et al.*, 2020). Studies have shown that fungal-bacterial interactions can promote plant growth and inhibit plant pathogens (Santoyo *et al.*, 2021; Artursson *et al.*, 2005). Therefore, the compatible interactions between *P. lilacinus* and *V. paradoxus* suggest their synergistic coexistence that may offer potential benefits and stress resilience in soybean plants

We further performed the split-root assay using sterile soil that provided valuable insights into the localized effects of enriched microbiomes on soybean growth. Plants (Clark) subjected to the SR3 treatment (microbiome/microbiome) exhibited slightly healthier shoots (non-significant) and healthier root systems along with the increased leaf chlorophyll content and rhizosphere siderophore compared to SR1 (control/control) and SR2 (control/microbiome). These findings emphasize the local effects of enriched microbiomes in promoting soybean growth. The involvement of local signal-promoting growth and disease resistance was also reported in sorghum (Kabir *et al.*, 2024) and tomato (Lucas *et al.*, 2018). However, studies also reported that systemic colonization by beneficial microbes may optimize physiological traits, including photosynthetic efficiency and biomass accumulation (Chi *et al.*, 2005; Harman *et al.*, 2021). In this study, the effects of *P. lilacinus* and *V. paradoxus* inoculum are context-dependent in soybean, with localized signals dominating in compartmentalized systems, while systemic colonization optimizes physiological traits under more uniform conditions.

Building on the compatibility of *P. lilacinus* and *V. paradoxus*, we validated their persistence and impact on sensitive genotypes to the combined Fe deficiency and drought stress. Microbial inoculants have gained significant attention for their potential to enhance plant tolerance to abiotic stresses (Gonçalves *et al.*, 2023; Khan *et al.*, 2023). Our study found that microbiomes enriched in Clark (tolerant) improved plant growth and stress resilience in genotypes sensitive to Fe deficiency and drought. Chen et al. (2023) reported that *P. lilacinus*, when applied in combination with *Bacillus subtilis*, enhanced watermelon growth by improving soil phosphorus availability and altering microbial community composition. Among plant symbionts, *Variovorax* is metabolically versatile and is known to modulate the production of indole-3-acetic acid (Sun *et al.*, 2018; Han *et al.*, 2011). Evidence from another study indicates that the ethylene signaling pathway plays a role in how *V. paradoxus* promotes leaf development and flowering in *Arabidopsis thaliana* (Chen *et al.*, 2013). Several studies have demonstrated the beneficial effects of diverse microbiomes in mitigating single abiotic stresses in plants, including but not limited to *Burkholderia phytofirmans* for drought in wheat (Naveed *et al.*, 2014), *Azospirillum brasilense* for drought in Arabidopsis (Cohen *et al.*, (2015), and *Trichoderma harzianum* for Fe deficiency in pea (Thapa *et al.*, 2025b). One of the key observations in our study was the effectiveness of the enriched microbiome in Arisoy, even when cultivated under combined Fe deficiency and drought, in nexus with elevated rhizosphere siderophore and root flavonoid levels. This suggests a synergistic interaction between the microbial inoculants, which form symbiotic associations with host plants, leading to stress mitigation in soybean under combined Fe deficiency and drought. However, AG58XF3 may possess metabolites other than flavonoids that facilitate symbiotic trade-offs with enriched microbiomes. Host plants release a diverse array of metabolites to facilitate and promote symbiotic interactions with microbial partners (Hacquard and Martin, 2024). Hence, it is essential to study the genetic interplay of plants and their microbiomes along with environmental heterogeneity. While our experiments were conducted under controlled greenhouse conditions, it is important to acknowledge that such environments do not fully capture the complexity and variability of field conditions. Therefore, future studies in natural field settings are necessary to validate the consistency and applicability of these findings in real-world agricultural systems.

**In conclusion,** this study embodies the first detailed investigation into the complex interplay of physiological, molecular, and microbial factors improving stress tolerance in soybean plants exposed to combined Fe deficiency and drought. The ability of Clark to combat stress is associated with nutrient and redox balance, along with Fe retention and physiological adjustment, possibly driven by elevated jasmonic and salicylic acid in the roots. The upregulation of symbiotic genes and increased levels of siderophores and root flavonoids in Clark were correlated with improved nutrient status and microbial enrichment, predominantly by *Variovorax* and *Paecilomyces* in the endosphere (Fig. 10). Furthermore, the compatibility and synergistic effects of enriched microbiomes, particularly in sensitive genotypes, highlight the potential of microbial inoculants to enhance stress resilience in soybean. Collectively, these findings offer actionable insights into agricultural innovation. The identified host traits and microbial partners serve as promising targets for breeding programs aimed at developing stress-tolerant soybean cultivars. Moreover, the beneficial microbes enriched in stress-resilient genotypes could be harnessed to formulate next-generation, microbiome-based biofertilizers tailored to improve plant health and productivity under challenging environmental conditions. By bridging plant genetics with microbial ecology, this work lays the foundation for translational strategies to enhance resilience not only in soybean but also in other legume crops facing abiotic stress under climate change scenarios.

**Fig. 10.**
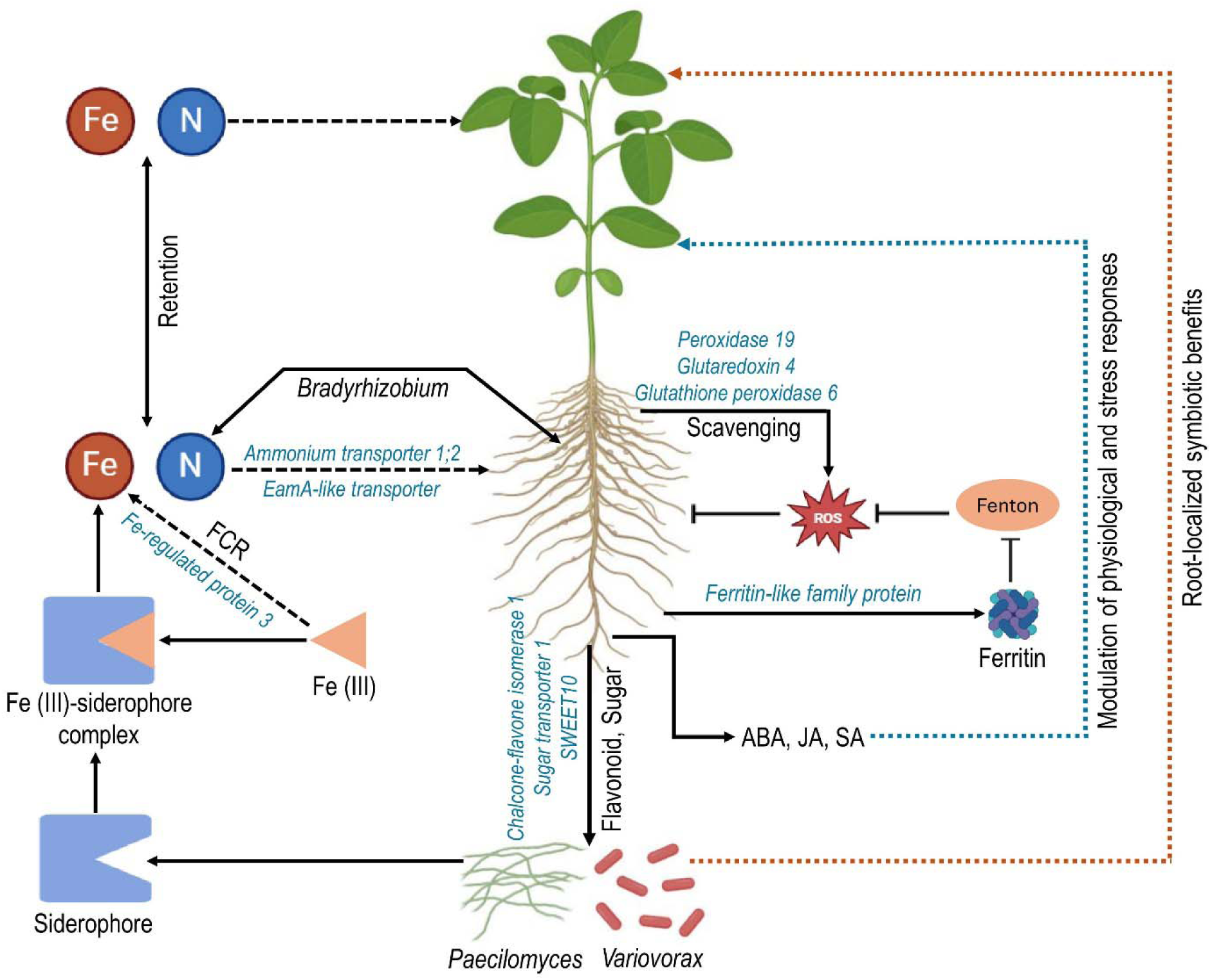
A model of stress resilience in Clark exposed to Fe deficiency and drought. The model illustrates the mechanisms underlying stress resilience, including enhanced Fe and N retention mediated by root-localized symbiotic interactions with *Bradyrhizobium*, *Paecilomyces*, and *Variovorax*. Key transporters such as *Ammonium transporter 1;2*, *EamA-like transporter*, *Sugar transporter 1*, and *SWEET10* facilitate nutrient exchange and carbon flow. Fe homeostasis is maintained through the activity of ferric chelate reductase (FCR), *Fe-regulated protein 3*, and siderophore-mediated Fe acquisition. Reactive oxygen species (ROS) are scavenged by *Peroxidase 19*, *Glutaredoxin 4*, and *Glutathione peroxidase 6*, while ferritin provides additional protection by inhibiting Fenton reactions. Flavonoids and sugars produced in the roots further support microbial recruitment and hormonal modulation involving ABA, JA, and SA, collectively enhancing physiological and stress responses. This integrated system highlights the synergy between microbial symbiosis and plant metabolic pathways in resilience to combined Fe deficiency and drought stress in soybean.

## Supplementary data

Supplementary Table S1. Physico-chemical properties of soil used for the cultivation of soybean.

Supplementary Table S2: Changes in growth parameters and photosynthetic attributes in Clark and Arisoy were grown on vermiculite under sterile conditions (without any microbes) for both the control and Fe-Drought+ conditions. Different letters indicate significant differences between the control and Fe-Drought+ conditions at a *p* <0.05 level, based on *t*-test. The data represents the means ± SD of three independent biological samples (*n* = 3).

Supplementary Table S3: List of all differentially expressed genes (DEGs) and raw read counts per gene generated using HTSeq in response to treatment in two soybean genotypes: Clark and Arisoy.

Supplementary Table S4: Changes in the expression of stress-related genes and their homologs in the roots Clark and Arisoy under Fe-Drought+ conditions. Asterisks (*) denote significant differences at *p* < 0.05. Positive values indicate upregulation, while negative values indicate downregulation. Data represents the average of three independent biological replicates.

Supplementary Table S5. List of primers used for qPCR experiments.

Supplementary Fig. S2. RNA-seq data quality: (A) average base quality scores across read positions for all sequenced samples. Each colored line represents an individual sample (e.g., 13_S114 to 24_S125). The x-axis indicates the base position within each read (1–150 bp), and the y-axis shows the corresponding average Phred quality score and (B) Total number of sequencing reads per sample. Bar graph representing the total raw reads obtained from each sample (13_S114 to 24_S125). The y-axis indicates the number of reads in millions (M), showing high and consistent sequencing depth across most samples, with all samples exceeding approximately 13 million reads.

Supplementary Fig. S2. Cross-sections of root nodules from Clark and Arisoy genotypes cultivated under control and Fe-Drought+ conditions.

Supplementary Fig. S3: Cultivation of pure *Paecilomyces lilacinus* and *Variovorax paradoxus* in nutrient broth.

## Acknowledgment

We express our gratitude to the Genomics Core of Michigan State University and CD Genomics. This research was supported by a startup grant (5SFAES-293007) and Elizabeth and Hayden Cutler’s Endowed Professorship in Biotechnology from the University of Louisiana at Monroe. We also thank Mr. Timothy McMahan for his assistance with Nanopore Sequencing.

## Author contributions

MRH was involved in experimental design, plant cultivation, RNA extraction, RNA-seq data analysis, interpretation, 16S data analysis, split-root assay, microbial co-culture experiments, and prepared the manuscript draft. AT participated in plant phenotyping, performed biochemical assays, and extracted nodule DNA. AHK provided overall supervision, guidance in experimental design and bioinformatics analytics, and manuscript revision.

## Data availability

The data supporting the findings of this study are available in the article and its supplementary information files. Also, the sequencing data used and described in this study has been uploaded to NCBI under the Bioproject accession numbers: PRJNA1204368 (RNA-seq in root) and PRJNA1204372 (16S in root), PRJNA1204386 (ITS in roots) and PRJNA1204446 (16S in nodule).

