## Supplementary for "Iron retention coupled with trade-offs in localized symbiotic effects confers tolerance to combined iron deficiency and drought in soybean"

**Supplementary Table S1.** Physico-chemical properties of soil used for the cultivation of soybean.

| **Texture**  Clay | | | | | **Organic matter**  6% | | | | | **Cation Exchange Capacity**  16.2 meq/100g | | | | **Water Holding Capacity**  44% | | | |
| --- | --- | --- | --- | --- | --- | --- | --- | --- | --- | --- | --- | --- | --- | --- | --- | --- | --- |
| **Elemental content ppm** | | | | | | | | | | | | | | | | | |
| Al aluminum | N arsenic | B boron | Ca calcium | Cd cadmium | Cr chromium | Cu copper | Fe iron | K potassium | Mg magnesium | Mn manganese | Mo molybdenum | Na sodium | Ni nickel | P phosphorus | Pb lead | S sulfur | Zn zinc |
| no limit | no limit | no limit | no limit | <2 | <100 | <100 | no limit | no limit | no limit | <3500 | <440 | no limit | <50 | no limit | <75 | no limit | <100 |
| 5352 | 121 | 11.21 | 1234 | 0.21 | 8.8 | 12.4 | 9102 | 1177 | 1687 | 763 | 0.48 | 887 | 8.46 | 659 | 17.2 | 177 | 71.5 |

**Supplementary Table S2:** Changes in growth parameters and photosynthetic attributes in Clark and Arisoy were grown on vermiculite under sterile conditions (without any microbes) for both the control and Fe-Drought+ conditions. Different letters indicate significant differences between the control and Fe-Drought+ conditions at a *p* <0.05 level, based on *t*-test. The data represents the means ± SD of three independent biological samples (*n* = 3).

| Growth parameters | Clark | | Arisoy | |
| --- | --- | --- | --- | --- |
|  | Control | Fe-Drought+ | Control | Fe-Drought+ |
| SPAD | 32.16±3.62^a^ | 14.33±1.07^b^ | 35.56±2.97^a^ | 16.16±2.4^b^ |
| Shoot height (cm) | 43.16±0.7^a^ | 30.46±1.12^b^ | 50.03±8.66^a^ | 26.3±3.63^b^ |
| Stem diameter (mm) | 2.36±0.28^a^ | 2.03±0.11^a^ | 2.03±0.15^a^ | 1.2±0.25^b^ |
| Number of leaves | 16.66±2.88^a^ | 10.66±1.52^a^ | 17.33±3.51^a^ | 7.66±0.57^b^ |
| Root length (cm) | 18.46±3.88^a^ | 12.23±1.1^a^ | 19.76±4.1^a^ | 5.56±1.67^b^ |
| Shoot fresh weight (g) | 6.58±0.19^a^ | 1.76±0.1^b^ | 3.66±0.7^a^ | 1.02±0.52^b^ |
| Shoot dry weight (g) | 1.51±0.23^a^ | 0.4±0.01^b^ | 0.85±0.21^a^ | 0.28±0.14^b^ |
| Leaf RWC (%) | 84.53±10.05^a^ | 76.08±5.46^b^ | 81.7±5.27^a^ | 57.45±8.05^b^ |

**Supplementary Table S3:** List of all differentially expressed genes (DEGs) and raw read counts per gene generated using HTSeq in response to treatment in two soybean genotypes: Clark and Arisoy. See uploaded Excel sheet.

**Supplementary Table S4:** Changes in the expression of stress-related genes and their homologs in the roots Clark and Arisoy under Fe-Drought+ conditions. Asterisks (^*^) denote significant differences at *p* < 0.05. Positive values indicate upregulation, while negative values indicate downregulation. Data represents the average of three independent biological replicates.

| Genes | *Arabidopsis thaliana* homolog and similarity % | Clark | Arisoy |
| --- | --- | --- | --- |
| *GLYMA.20G241500 (Chalcone-flavone isomerase 1)* | Chalcone-flavanone isomerase family protein: 63% | 2.7^*^ | -2.5 |
| *GLYMA.17G027400 (Drought-induced-19 protein)* | Late embryogenesis abundant 3: 68% | 3.8^*^ | 3.6^*^ |
| *GLYMA.11G106900 (Dehydration-associated protein)* | Senescence/dehydration-associated protein-like protein: 76.5% | 2.4^*^ | 1.5 |
| *GLYMA.07G032600 (Glutaredoxin 4)* | Glutaredoxin 4: 86.2% | 2.9^*^ | -0.04 |
| *GLYMA.05G035500 (EamA-like transporter)* | Cationic amino acid transporter 2: 100% | 2.7^*^ | 0.0 |
| *GLYMA.08G013800 (Glutathione peroxidase 6)* | Glutathione peroxidase 6: 82.2% | 2.6^*^ | 0.0 |
| *GLYMA.10G167800 (Ammonium transporter 1;2)* | Ammonium transporter 1;2: 86.2% | 3.1^*^ | -3.41 |
| *GLYMA.10G146600 (Iron-regulated protein 3)* | Iron-regulated protein 3: 92.1% | 2.7^*^ | -2.0 |
| *GLYMA.13G172300 (Peroxidase 19)* | Peroxidase superfamily protein: 97.1% | 2.3^*^ | -0.73 |
| *GLYMA.15G154100 (2-oxoglutarate)* | 2-oxoglutarate: 90% | 2.7^*^ | 0.12 |
| *GLYMA.07G113500 (ATPase 11)* | H^+^-ATPase 4: 100% | 2.8^*^ | 4.1^*^ |
| *GLYMA.08G059700 (Sugar transporter 1)* | Sugar transporter protein 12: 85% | 2.9^*^ | 2.1 |
| *GLYMA.07G006500 (Sulfate transporter 3;4)* | Sulfate transporter 3;1: 100% | 2.5^*^ | -4.3^*^ |
| *GLYMA.19G091400 (Ferritin-like family protein)* | Ferritin/ribonucleotide reductase-like family protein: 88.3% | 2.4^*^ | 1.8 |
| *GLYMA.17G119400 (Stabilizer of iron transporter SufD)* | Stabilizer of iron transporter SufD: 100% | 3.3^*^ | 1.1 |
| *GLYMA.04G198400 (Bidirectional sugar transporter SWEET10)* | Nitrate transporter 1.5: 100% | 3.1^*^ | 0.2 |

**Supplementary Table S5.** List of primers used for qPCR experiments.

| **Genes** | **Primers** |
| --- | --- |
| *GAPDH* | Forward: TCCTTTCATTACCACCGATTACA  Reverse: CGATCTGAACCTATCCGAACAA  Amplicon size: 105 bp |
| *Tubulin* | Forward: GCTTGATAATGAGGCGCTCTA  Reverse: CTCATGGTTGCGGAGATCAA  Amplicon size: 99 bp |
| *GLYMA.20G241500 (Chalcone-flavone isomerase 1)* | Forward: CCGCTAAGTGGAAGGGTAAA  Reverse: TGGAGAAAGTAGGTAGGTAGGT  Amplicon size: 98 bp |
| *GLYMA.04G198400 (Bidirectional sugar transporter SWEET10)* | Forward: TGTGGCTTGGCGTTTCTAT  Reverse: TGATGCAGCTCCCAAATACTC  Amplicon size: 85 bp |
| *GLYMA.19G091400 (Ferritin-like family protein)* | Forward: GTGGATATTGAGCGGGAGTT  Reverse: CGATCTGCCACAAACTCAATG  Amplicon size: 101 bp |
| *GLYMA.10G146600 (Iron-regulated protein 3)* | Forward: GGCAACCGACTATGCAATTAAAG  Reverse: AACTCCCACCCACCCTAATA  Amplicon size: 98 bp |
| *GLYMA.17G027400 (Drought-induced-19 protein)* | Forward: GTTCTCTCTCGCAAGCCAAA  Reverse: CTGACCGATGAACGGGAATTAG  Amplicon size: 75 bp |

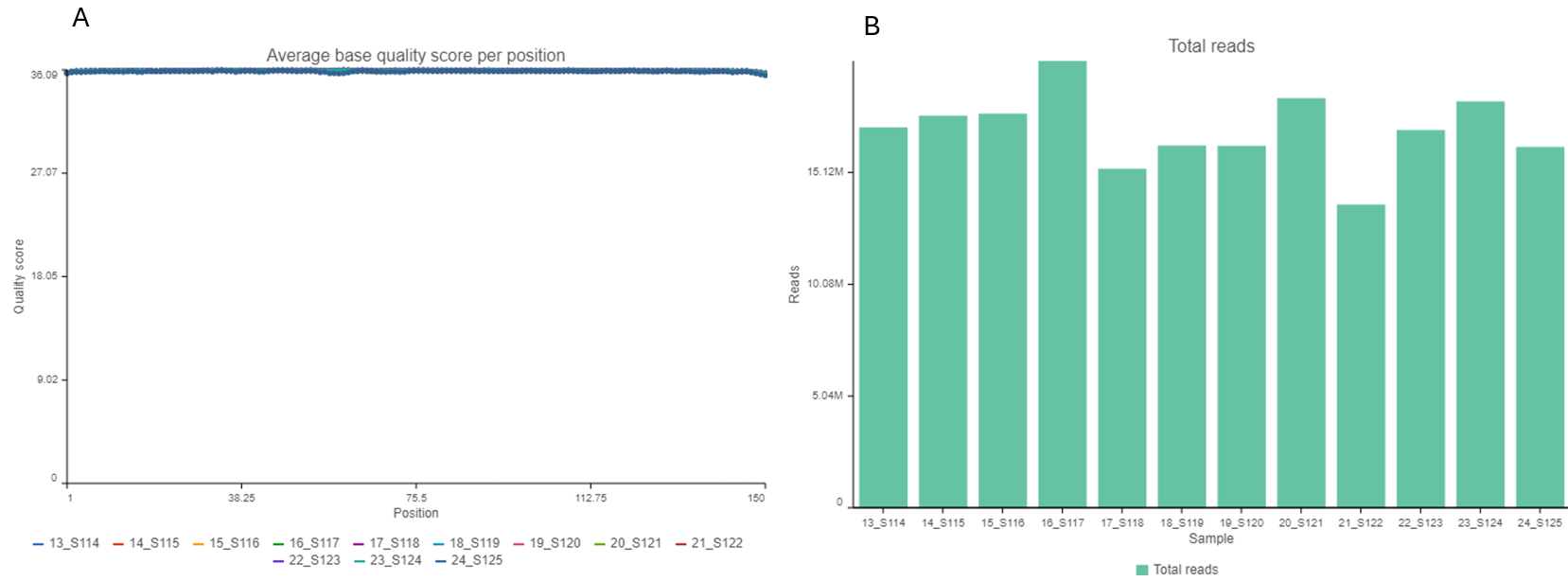

**Supplementary Fig. S2.** RNA-seq data quality: (A) average base quality scores across read positions for all sequenced samples. Each colored line represents an individual sample (e.g., 13_S114 to 24_S125). The x-axis indicates the base position within each read (1–150 bp), and the y-axis shows the corresponding average Phred quality score and (B) Total number of sequencing reads per sample. Bar graph representing the total raw reads obtained from each sample (13_S114 to 24_S125). The y-axis indicates the number of reads in millions (M), showing high and consistent sequencing depth across most samples, with all samples exceeding approximately 13 million reads.

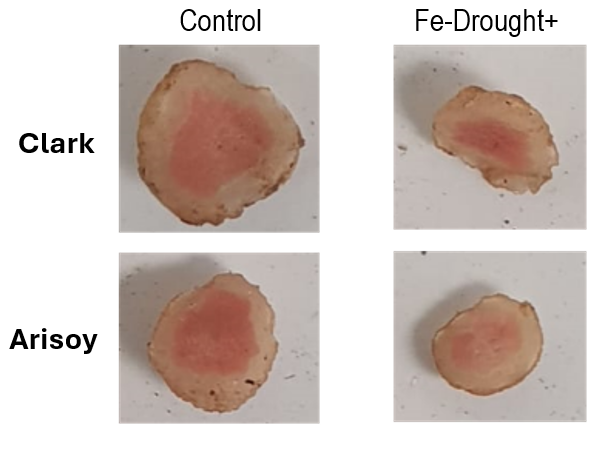

**Supplementary Fig. S2.** Cross-sections of root nodules from Clark and Arisoy genotypes cultivated under control and Fe-Drought+ conditions.

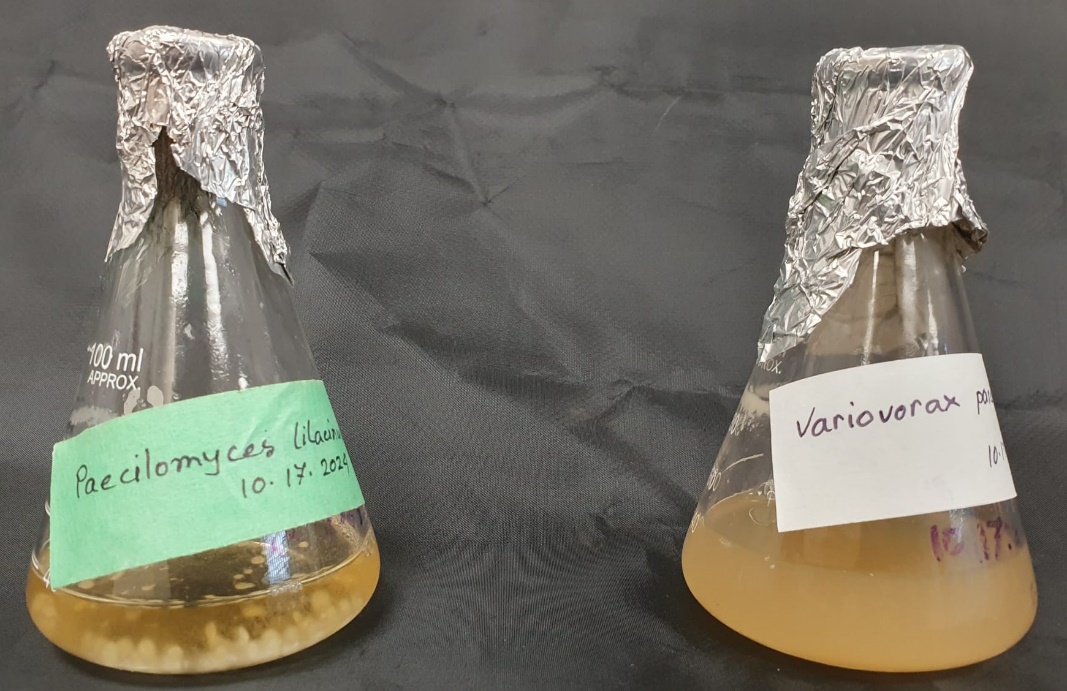

**Supplementary Fig. S3:** Cultivation of pure *Paecilomyces lilacinus* and *Variovorax paradoxus* in nutrient broth.
